# Direct Observation of a Coil-to-Helix Contraction Triggered by Vinculin Binding to Talin

**DOI:** 10.1101/741884

**Authors:** Rafael Tapia-Rojo, Alvaro Alonso-Caballero, Julio M. Fernandez

## Abstract

Vinculin binds unfolded talin domains in focal adhesions, which recruits actin filaments to reinforce the mechanical coupling of this organelle. However, the mechanism by which this interaction is regulated, and its impact in the force transmission properties of this mechanotransduction pathway remain unknown. Here, we use magnetic tweezers force spectroscopy to measure the binding of vinculin head to the talin R3 domain under physiological force loads. For the first time, we resolve individual binding events as a short contraction of the unfolded talin polypeptide due to the reformation of the helices in the vinculin-binding sites. This force-dependent contraction dictates the mechanism by which force regulates the talin-vinculin interaction. Force is needed to unfold talin and expose the cryptic vinculin-binding sites; however, the structural contraction triggered by binding introduces an energy penalty that increases with force, defining an optimal binding force range. This novel mechanism implies that the talin-vinculin-actin association works in focal adhesions as a negative feedback mechanism, which operates to stabilize the force acting on each junction.

## I. INTRODUCTION

Cell function relies largely on the ability of cells to interpret their mechanical environment, and respond dynamically to these force cues—mechanotransduction [1–4]. Cells anchor the extracellular matrix and probe its stiffness through focal adhesions, which connect transmembrane integrins with the active cellular cytoskeleton and regulate the transmission and transduction of force into biochemical regulatory signals [5–7]. The interaction between integrins and F-actin filaments is done through adaptor proteins like talin, that establish this physical connection but also regulate the mechanical response of this organelle [8–10]. Talin binds integrin cytodomains through its N-terminal FERM head, which is followed by a flexible rod region formed by 13 helical bundle domains (Fig. S1) [11]. The talin rod is the mechanosensitive region and responds to force by establishing a complex network of interactions with several other molecular partners, whose recruitment depends on the mechanical cue on talin [12–14]. Among them, vinculin is particularly relevant, as the 11 vinculin-binding sites distributed along the talin rod are cryptic, and require mechanical unfolding of its helical domains for vinculin to bind [11, 15, 16]. Upon binding, vinculin recruits F-actin filaments, which reinforce the mechanical coupling and increase the strength of the focal adhesion [17–20]. This mechanism has been suggested to operate as a positive feedback; as the force across talin increases gradually, its domains unfold and more vinculin molecules bind, increasing actin recruitment and subsequent force transmission [21–23]. However, vinculin is also required for the stabilization of adhesions under force [24]; hence, it remains unknown how vinculin binding could regulate force application and control the lifetime of focal adhesions. Each vinculin-actin linkage bears around 2.5 pN of force that contributes to the overall tension of the linkage [24], but vinculin dissociates from talin under excessive force loads [25]. This could suggest that vinculin binding occurs on a restricted force regime over which each linkage should operate. However, the force-dependency of the talin-vinculin interaction has never been measured, and the mechanism by which force regulates this complex remains unknown.

Here, we use magnetic tweezers force spectroscopy to measure binding of vinculin headto the talin R3 domain and investigate how force modulates this interaction. Thanks to the improved resolution of our custom-made setup, we resolve for the first time individual vinculin head binding events as a short contraction of the talin polypeptide, due to a coil-to-helix transition induced by binding (Fig. 1). The scaling of this contraction with force indicates that vinculin binding is cooperative, and two vinculin head molecules bind simultaneously to the unfolded R3 domain. Our experiments reveal a biphasic force dependency of the binding reaction. First, force favors binding by unfolding talin and exposing the cryptic binding sites. However, for vinculin head to bind, it must do mechanical work against the force to contract the stretched talin polypeptide, which is unfavorable at higher forces. This novel mechanism regulates talin-vinculin interaction and defines an optimal force range for binding. By integrating our findings into a minimalistic model, we demonstrate that the talin-vinculin-actin association might operate in focal adhesions as a negative feedback mechanism, which recruits or dissociates vinculin molecules to stabilize the force on each junction to an optimal value.

**FIG. 1.**
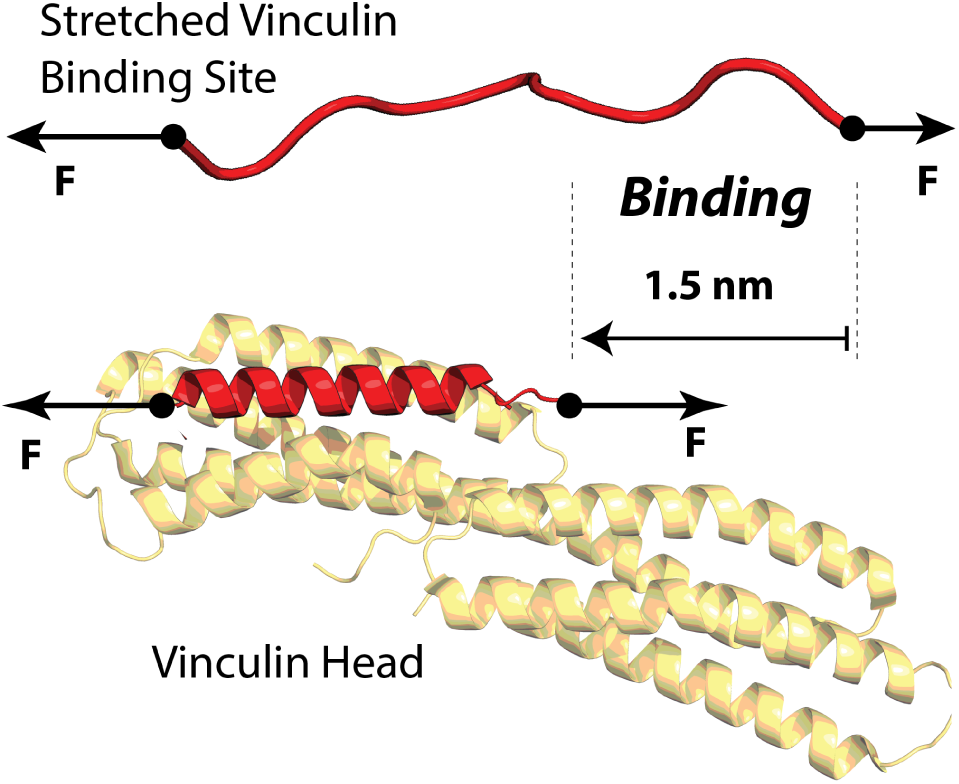
Vinculin binding requires the structural accommodation of the talin polypeptide on the vinculin head: Under force, talin unfolds and the vinculin-binding sites become unstructured polypeptide chains. Upon binding of one vinculin head molecule, the binding site helix must reform, which shortens talin by ∼1.5 nm at a force of 9 pN.

## II. RESULTS

### A. Real Time Detection of Vinculin Binding to Talin

To isolate individual vinculin binding events, we study the talin R3 domain, a four-helix bundle located in the N-terminal region of the talin rod (Fig. S1) [11]. This domain has two vinculin-binding sites and remarkable low stability due to the four-threonine belt buried in its hydrophobic core [11, 22, 23]; hence, it is ideally placed to play a key role in talin activation by force, by recruiting a cluster of vinculin molecules at low forces, which would amplify the mechanical coupling, and contribute to the maturation of the focal adhesion. Mutation of these four threonines to hydrophobic valine and isoleucine residues increases the domain stability while leaving its vinculin-binding properties intact [11].

Here, we study vinculin binding both on the R3 WT and R3 IVVI but focus the characterization of the mutant since its higher mechanical stability amplifies the mechanical signature for vinculin binding; however, the binding mechanism is completely equivalent on both domains. We use our custom-made magnetic tweezers to apply physiological forces to single R3 domains in the presence of vinculin head, and measure its extension changes in real-time with nm resolution. Our molecular construct contains the R3 domain (either the WT or IVVI mutant) followed by eight repeats of titin I91 domain as molecular handles, flanked by a HaloTag enzyme for covalent tethering to the glass surface, and biotin for anchoring to a streptavidin-coated superparamagnetic bead (Fig. 2A; see SI for methods). Forces between 0.1 and 120 pN are applied with a sub-pN resolution by generating a magnetic field with a pair of permanent magnets, or a magnetic tape head [26, 27].

**FIG. 2.**
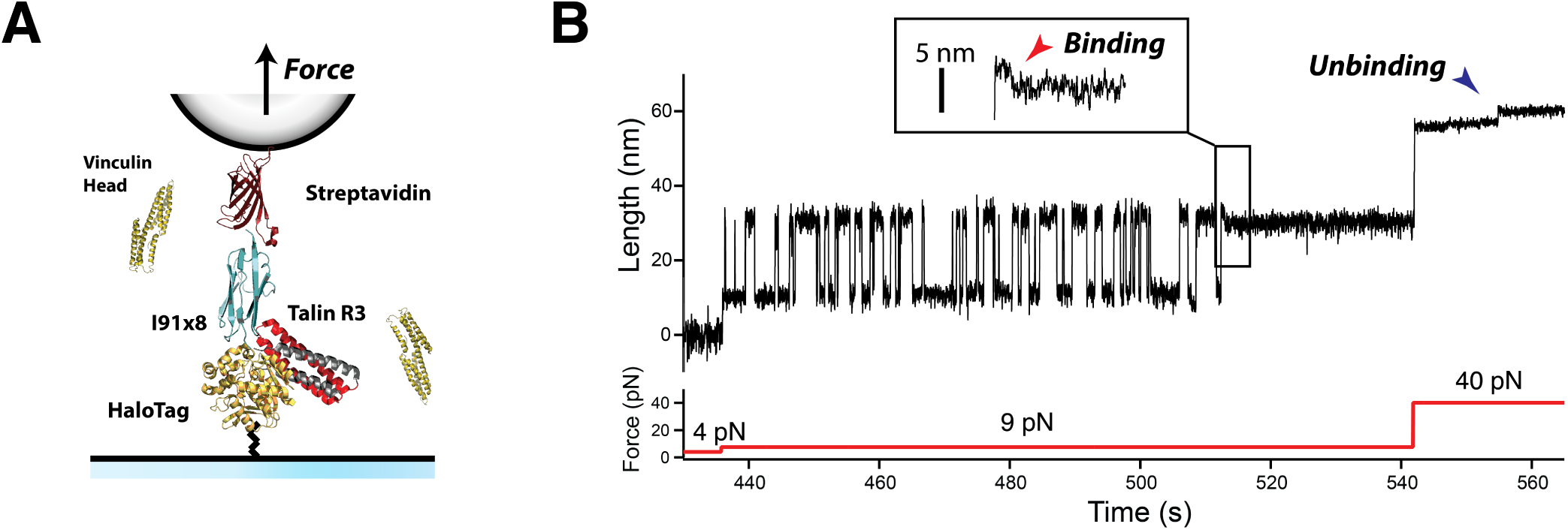
Real-time detection of the coil-to-helix contraction induced by vinculin head binding: (A) Schematics of a magnetic tweezers experiment for detecting vinculin binding events. We engineer a (R3)-(I91)_8_ protein construct, flanked by a HaloTag, for covalent tethering to a glass coverslip, and a biotin, for anchoring to streptavidin-coated superparamagnetic beads. Physiological-level forces in the pN range are applied through a magnetic field gradient created either by a pair of permanent magnets or a magnetic head. The experiment is conducted in the presence of vinculin head, and the extension changes due to folding or binding are measured with nm resolution. (B) Magnetic tweezers recording showing individual vinculin head binding events to the R3 IVVI domain. R3 IVVI folds and unfolds in equilibrium, which yields extension changes of ∼20 nm. In the presence of 20 nM vinculin head, these dynamics eventually stop due to binding of vinculin head. This event is resolved as a ∼3 nm contraction that occurs in the unfolded talin polypeptide due to the reformation of the *α*-helices of its two vinculin-binding sites (red arrow, inset). The complex dissociates at higher forces, showing ∼3 nm upward steps (blue arrow).

Figure 2B shows our single-molecule assay for vinculin head binding detection. Starting from folded R3 IVVI at 4 pN, we increase to 9 pN, where the R3 IVVI domain exhibits reversible folding/unfolding dynamics (see [27], and Fig. S2 for the mechanical characterization of this R3 IVVI and R3 WT). The unfolding/folding transitions are resolved as ascending/descending ∼20 nm changes in the extension of R3 due to the transition between the folded state and an unstructured polymer. Previous force spectroscopy studies showed that vinculin head binding blocks talin refolding [25]. Our experiments confirm this observation; in the presence of 20 nM vinculin head, R3 folding dynamics cease after a few seconds, and the protein is locked on its unfolded state (Fig. 2B). This blocked state extends for several hours, in contrast with the unaltered R3 folding dynamics that show in the absence of vinculin head (Fig. S3). The improved resolution of our instrument and the use of a constant force, allows us to observe the binding event as a short contraction of the unfolded polypeptide that always precedes the arrest of R3 folding dynamics (Fig. 2B; inset, red arrow). This contraction indicates that vinculin head binding induces a conformational change on unfolded R3, likely a coil-to-helix transition required for vinculin head to firmly bind its substrate (Fig. 1). This bound state can be reversed by a high force pulse, where the opposite transition occurs, and ∼3 nm upward steps are observed (blue arrow), after which the R3 domain recovers its ability to fold (Fig. S4). Experiments with R3 WT in presence of vinculin head reveal this same effect; however, at the coexistence force for R3 WT (5 pN), the binding contraction is too small to be detected, and is only observed at forces >8 pN (Fig. S5).

### B. Cooperative Binding of Vinculin Head to the Talin R3 Domain

Our measurements demonstrate that the mechanical fingerprint for vinculin head binding is a contraction of few nanometers on the unfolded R3 polypeptide. Structural studies have determined that vinculin-binding sites are amphipathic six-turn *α*-helices, buried in the core of talin domains by extensive hydrophobic interactions [28]. Upon binding, this helix is inserted intimately into vinculin head, which displaces the initial interaction between the head and tail of vinculin, present in its autoinhibited state [29]. Hence, this helical structure is required for vinculin recognition and binding, which might explain the need for the structural contraction we observe upon binding.

Figure 3A shows averaged recordings of individual vinculin head binding events on R3 IVVI, measured at different forces. The size of the contraction induced by binding increases with force, which is an expected observation in the transition from a random-coiled chain to a compact structure. Interestingly, in all our experiments, we observe a single contraction event, sufficient to form the bound state even though the R3 domain has two vinculin-binding sites. By contrast, when dissociating mechanically at forces above 40 pN, we resolve in most cases two distinct steps with an extension of ∼3 nm (Fig. 3B). In some traces, we observe only a single unbinding step, likely because the first one occurred too fast to be resolved since two unbinding steps are required for complete dissociation (Fig. S4). An analysis of the unbinding kinetics confirms that a fraction of unbinding events occurs within the resolution limit of our instrument (Fig. S6).

**FIG. 3.**
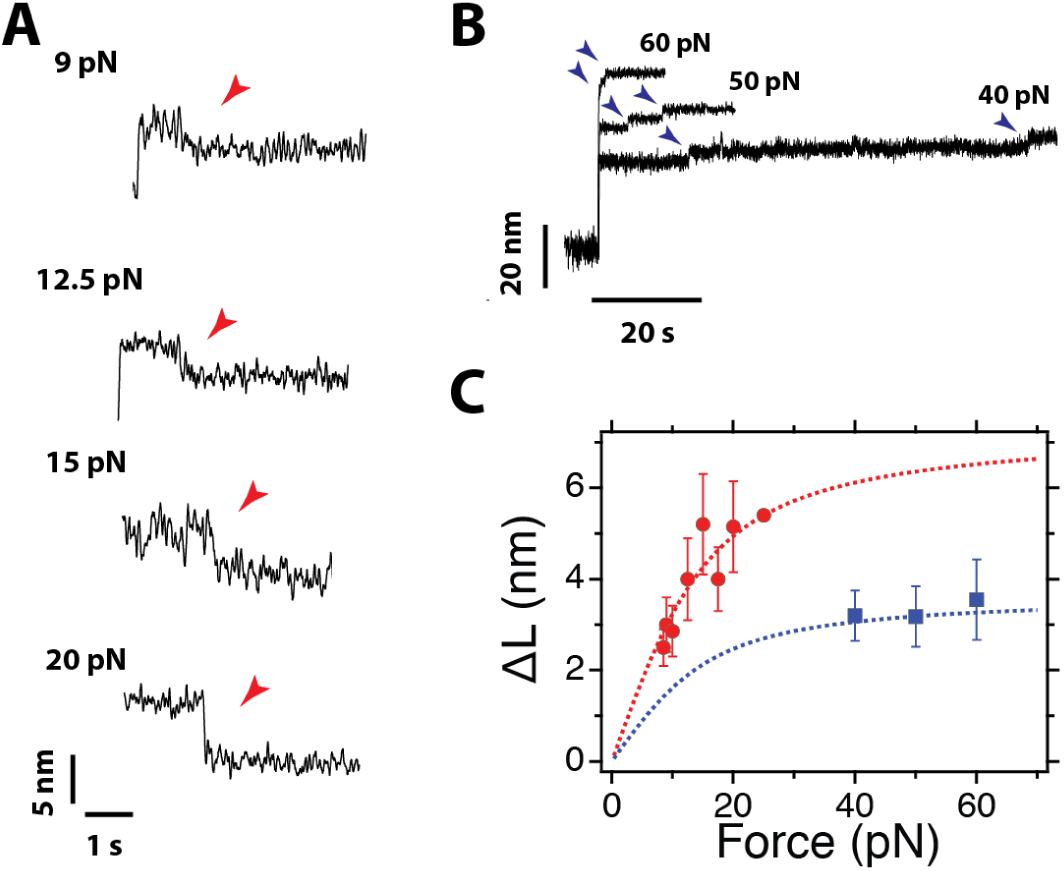
Vinculin binding contracts unfolded talin: (A) Averaged recordings of the binding contractions at different pulling forces. The magnitude and duration of the contraction depend on the force. Traces averaged from >10 recordings. (B) Unbinding steps at different pulling forces. Two ∼3 nm steps are observed, after which talin recovers its ability to refold. (C) Average step sizes for the binding contractions (red) and unbinding steps (blue) measured as a function of the pulling force. The binding contraction scales with force following the FJC polymer model with a contour length of 7.3 nm, which agrees with the simultaneous formation of the two *α* helices of the vinculin-binding sites. The steps of unbinding have half that contour length, indicating that they correspond to the unraveling of a single binding site helix. Error bars are the SEM; data collected over 35 molecules, 156 binding steps, and 501 unbinding steps.

Our data suggest that two vinculin head molecules bind simultaneously to unfolded R3, while each vinculin head unbinds independently. The magnitude of the binding and unbinding events scales with force following the freely-jointed chain (FJC) model for polymer elasticity [30]. We measure the size of the binding contraction as a function of force (Fig. 3C, red marks), and fit its dependence to the FJC model, obtaining a contour length change of Δ*L*_c_ = 7.33±0.69 nm (red dashed line). Plotting the same FJC fit with half that contour length results in a complete agreement with the steps measured for unbinding (Fig. 3C; blue). This indicates that the R3 sequence sequestered by each binding contraction is twice that liberated by each unbinding step. Given that each vinculin-binding site contains 19 residues, and that the extension of the formed helix is about 3.5 nm [11, 29], the expected contour length change for each coil-to-helix transition is 3.7 nm, in agreement with our measurements. An equivalent force scaling is measured in the binding/unbinding events on the R3 WT domain, which indicates that the structural transition triggered by binding is analogous both in the mutant and R3 WT domains (Fig. S5). Together, our observations confirm that each binding contraction corresponds to the simultaneous reformation of two vinculin-binding helices, while each unbinding step is the uncoiling of a single helix.

From this evidence, it remains uncertain which is the binding pathway followed by vinculin head to reach the bound state. One of the possible scenarios could involve repeated fluctuations between the coil and helix states in talin, with a first-hitting binding mechanism upon encounter of the appropriate substrate conformation. However, an interesting observation from our single-molecule recordings is that binding is not instantaneous but occurs as a slow relaxation, which, at 9 pN, takes as long as 500 ms, and that accelerates with force (Fig. S7). This suggests that vinculin head binding requires a maturation process, perhaps initiated by a recognition of key residues on the unfolded R3 polypeptide, followed by a sequential contraction and reformation of the helices. However, the rationale for the force-dependence observed in this maturation process remains inconclusive.

The simultaneous nature of the vinculin head binding event suggests a cooperative mechanism that should introduce a non-linear concentration dependence on the binding kinetics. From the perspective of single-molecule enzyme kinetics [31, 32], this process can be modeled as:

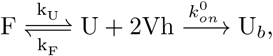

where F stands for folded R3, U for unfolded R3, U_b_ for the bound state, and Vh for the vinculin head domain. This process is governed by three kinetic rates: *k*_U_ and *k*_F_ are the unfolding and folding rates of R3, and 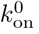 is the pseudo-first-order rate constant, which depends on the concentration of vinculin as 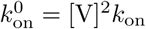, since two molecules are required to acquire the bound state. Experimentally, we measure the waiting time (*t*_b_) to observe the talin-bound state (Fig. 3A), and the binding rate *k*_b_ = 1*/t*_b_ can be derived analytically as (see SI, section IX):

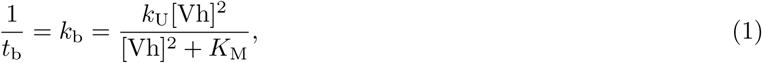

where *K*_M_ = (*k*_U_ + *k*_F_)*/k*_on_. This expression is the single-molecule analogous to a second-order Hill equation. Interestingly, in this case, the unfolding rate of talin plays the role of the maximum velocity of the reaction.

Figure 4A shows binding trajectories to R3 IVVI at 9 pN and different vinculin head concentrations. The waiting time *t*_b_ is determined from each recording as the time from the beginning of the 9 pN probe pulse, until the contraction event after which R3 folding dynamics stop. Figure 3B shows the distributions of waiting times at three representative concentrations, fitted to the expression derived from the kinetic model (see SI, section IX). Vinculin head binding kinetics are governed by two competing timescales, the folding/unfolding dynamics of R3, and the concentration-dependent on-rate 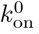. At low concentrations, the on-rate is much slower than R3 folding kinetics, and the process is rate-limited by vinculin head association. However, as the concentration increases, both opposing processes become comparable, and we observe a peaked distribution; binding cannot occur faster than the R3 domain unfolds. Unfortunately, it is not possible to do experiments at higher vinculin head concentrations, since binding occurs so fast that the fingerprint for vinculin head binding is lost.

**FIG. 4.**
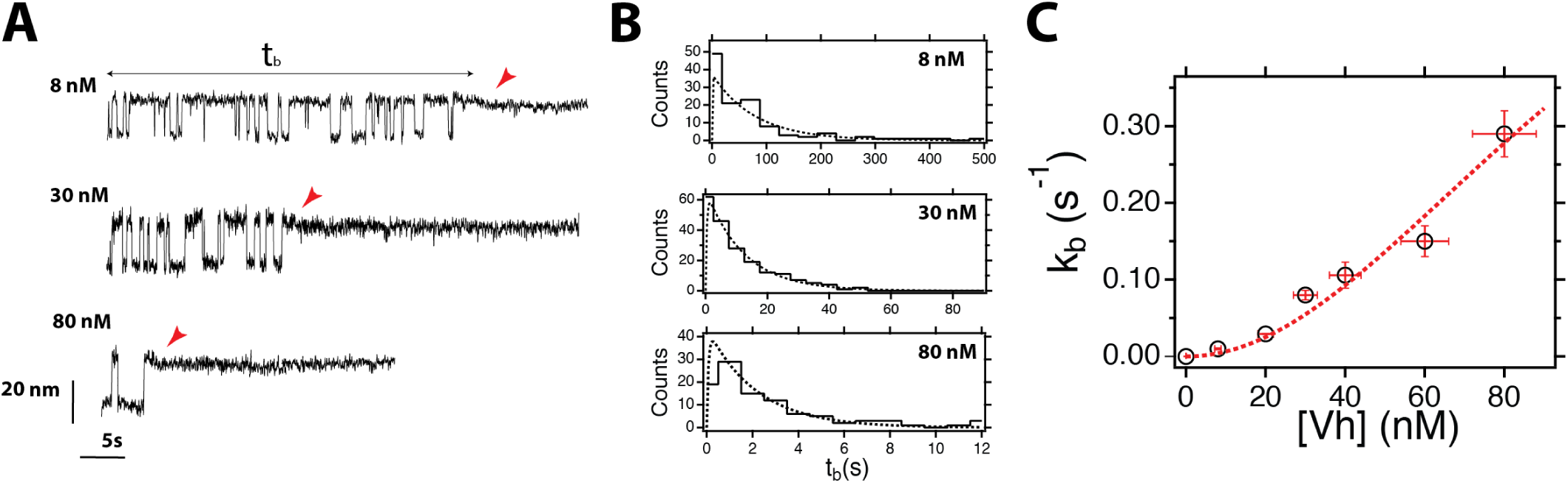
Stoichiometry of vinculin binding to talin: (A) Magnetic tweezers recordings of talin in presence of 8, 20, and 40 nM vinculin at a force of 9 pN. The waiting time for vinculin binding (*t*_b_) can be measured at the single-molecule level as the time taken since the probe force is set until the contraction is observed and the hopping dynamics stop. (B) Distribution of vinculin binding times at different concentrations, and at a force of 9 pN. The shape of the distribution follows a single-molecule enzymatic model, where two vinculin molecules bind simultaneously to unfolded talin. (C) Concentration dependence of the rate of vinculin binding, which follows a second-order Hill-like equation that demonstrates that two vinculin molecules bind simultaneously to the unfolded talin R3 domain. The point at 0 nM was estimated from three very long recordings in the absence of vinculin (5, 7, and 36 hours), that showed no arrest of binding dynamics. Vertical error bars are the SEM, and horizontal error bars are the precision on the determination of the vinculin head concentration (12.3 %). Number of events measured: 8 nM, N=122; 20 nM, N=184; 30 nM, N=200; 40 nM, N=267; 60 nM, N=139; 80 nM, N=104.

From these distributions, we calculate the binding rate *k*_b_, which has a quadratic dependence with the concentration, as described by Hill equation (Fig. 3C). This nonlinearity demonstrates the cooperative character of the vinculin head binding reaction. Fitting our data to Eq. 1 we obtain *k*_U_ = 0.54 ± 0.11 s^−1^, and *K*_M_ = 6852 ± 1480 nM^2^. The value of the unfolding rate agrees with that measured from R3 IVVI folding trajectories (Fig. S2). From *K*_M_, we obtain that half the maximum binding velocity occurs at ∼83 nM. Vinculin head has a strong tendency to aggregate, which might lead to uncertainties in the determination of its concentration (see Fig. S8).

Our observations are in agreement with previous evidence that suggested an affinity in the nanomolar range [29, 33]. Those experiments reported values between 3 and 30 nM, but were calculated using biochemical assays where vinculin **head** was left to interact with isolated vinculin-binding sites helices. However, the physiological affinity of vinculin to talin must be understood as a force-dependent quantity, which, as we demonstrated here, triggers structural changes on the binding substrate that should depend strongly on the tension applied to talin.

### C. Mechanical Force Regulates Vinculin Head Binding

Force is an essential actor in the interaction between vinculin and talin, being required to unfold talin domains and expose its cryptic sites. Additionally, and as previously reported [25], we have shown that **vinculin head** dissociates at high forces (>40 pN). This process arises likely from the destabilization of the reformed helices with force, which eventually will uncoil, and expel vinculin head. In this same sense, the binding reaction should be hampered by force, as the helices reform and contract a polymer that is mechanically stretched. This suggests that force could play a biphasic role in vinculin binding, first by establishing the threshold for talin unfolding, but also by hindering the coil-to-helix contraction as force increases.

We measure vinculin head binding to R3 IVVI at different forces and over a fixed time-window of 50 s, which readily demonstrates the biphasic effect of force on binding (Fig. 5A). At 8 pN, R3 IVVI explores the unfolded state with low probability, and vinculin head binds with slow kinetics. As force is increased to 9 and 10 pN, R3 IVVI unfolds more frequently, which results in faster vinculin head binding. However, at 15 and 20 pN, although R3 IVVI is always unfolded, the bindings kinetics rapidly slow down with force, until binding is blocked above 30 pN.

**FIG. 5.**
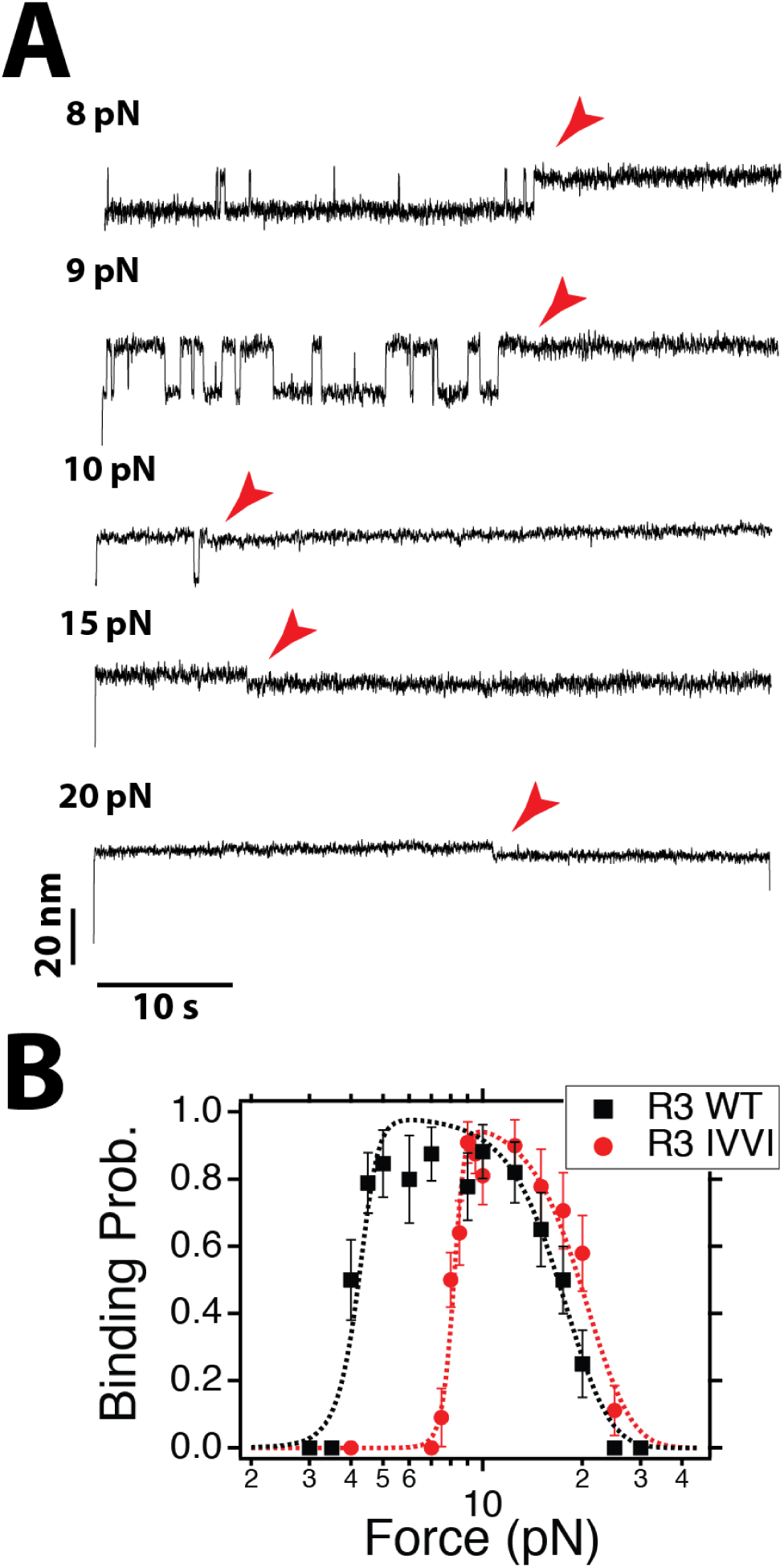
Mechanical force regulates vinculin head binding: (A) Typical recordings of vinculin binding at different forces during a 50 s time window and at a concentration of 20 nM. (B) Binding probability measured over a 50 s time window as a function of force. Force has a biphasic effect on binding, favoring it first by unfolding talin, but hampering it due to the energy penalty of the coil-to-helix contraction. The data is described by a simple model based on this mechanism (dashed line, see SI section X). Errors are SEM; data collected over 20 molecules and 259 observations for R3 IVVI, and 10 molecules and 213 observations for R3 WT.

This negative effect of force on binding arises from the coil-to-helix contraction that is triggered by binding. The R3 polypeptide shortens out of the equilibrium extension imposed by force; hence, vinculin binding does mechanical work against the pulling force, and this energy penalty increases steeply with force. To demonstrate the proposed mechanism, we measure the binding probability over a 50 s time-window (Fig. 5B), both on R3 WT and R3 IVVI. R3 WT has lower mechanical stability, showing equilibrium transitions between 4 and 6 pN, while R3 IVVI folds between 8 and 10 pN (Fig. S2). This difference in mechanical stability results in a lower threshold force for binding for R3 WT, compared to R3 IVVI. The binding probability quickly increases from 4 pN and saturates at 5 pN due to the sharp dependence of the unfolding rates (black squares), while for R3 IVVI the same behavior is observed at a higher force of 8 pN (red circles). However, the inhibitory effect of force arising from the coil-to-helix contraction is analogous for both domains, since the polymer properties of their binding sites are equal in both cases; the binding probability drops in the same fashion until binding is blocked at forces above 30 pN.

The mechanical work of binding can be estimated on a first approximation as Δ*W* ≈ *F* · Δ*L*(*F*), where Δ*L* is the force-dependent contraction measured in Fig. 2C. In this regard, we assume that the on-rate depends on force as 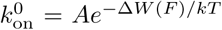. From this simple relation, we derive an analytical expression for the binding probability that incorporates the double effect of force on binding, controlled positively by the domain-dependent unfolding rate *k*_U_, but negatively by the on-rate 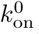 (see SI, section X for derivation). We use this expression to describe accurately our experimental data using only two free parameters (Fig. 5B, dotted lines; see Table S1 for parameter list). The agreement between the experimental data and our analytical description confirms the proposed mechanism for the mechanical regulation of the interaction between vinculin head and talin rod domains. Vinculin binding requires a coil-to-helix reformation, which is hampered by force. This, together with the force-induced exposure of the cryptic binding sites, defines the force regime over which vinculin binds. This mechanism can be directly extrapolated to other talin domains, or to ligand binding reactions that occur under similar conditions. The hierarchical mechanical stability of the talin rod domains will define a range of threshold forces for binding; however, the negative force dependency arises from the entropic penalty of the coil-to-helix transition required for binding, and, thus, can be expected to operate in a similar way for all vinculin-binding talin domains.

## III. DISCUSSION

Over the last 30 years, there has been an emphasis on understanding how molecular bonds respond to mechanical forces, a ubiquitous problem in biology. In his seminal 1978 paper, George Bell set the physical basis for the simplest case scenario; mechanical forces tilt the energy landscape in the pulling direction, decreasing linearly the height of the barrier, which results in an exponential decrease of the bond lifetime [34]. This simple theory—and more elaborated analytical corrections that followed—is used as a standard tool for analyzing the lifetime of biological bonds subject to pulling forces [35], and even other biological transitions, such as protein unfolding [36] or force-dependent chemical reactions [37]. In time, more complicated force dependencies have been measured, such as catch bond-like behaviors in the adhesive pili of some bacteria [38], or in the interaction between integrins and fibronectin [39]. However, how molecular interactions behave when force is applied to one of the components of the complex instead of to the bond itself remains poorly understood. This situation is of great generality and appears in physiological processes as diverse as DNA-protein interactions [40], antibody-antigen binding [41], or protein-protein interactions in cellular junctions [21], to name a few.

Here, we have proposed a molecular mechanism by which force regulates the formation of the mechanosensing complex between vinculin head and talin rod domains. Vinculin binding requires force to uncover its cryptic binding sites in talin, but force hampers this interaction as binding induces a structural rearrangement on the talin polypeptide. This double effect of force establishes an optimal binding force range that could define the mechanical regime over which cell adhesions operate. To explore this question, we propose a minimalistic model for the talin-mediated mechanical coupling between integrins and F-actin, which accounts for the talin-vinculin-actin association. This simple model integrates our measurements with the increased force transmission that occurs upon F-actin recruitment by vinculin, and predicts that the force-dependent interaction between talin and vinculin defines a negative feedback mechanism that stabilizes force across this linkage (see SI, section XII). In our simple scheme, an initial force input born by talin triggers unfolding of its bundle domains, promoting vinculin binding. The effect of vinculin is the recruitment of F-actin filaments, that elevate the force level on this junction, and subsequently increase further talin unfolding and vinculin binding. However, if the force on the linkage increases too much, vinculin binding becomes unfavorable, and the formed talin-vinculin complex dissociates, decreasing the overall force on the system. Hence, the force-dependence of the talin-vinculin interaction defines a mechanical negative feedback mechanism, which could explain how vinculin recruitment stabilizes force on focal adhesions [24].

We run Monte Carlo simulations on our model, built by concatenating 10 identical talin domains with the properties we measured for R3 IVVI (Fig. S9). The simulations are initiated with an arbitrary force input, which could arise from the mechanical coupling with the extracellular matrix, or the initial actin recruitment by talin. Upon unfolding of a talin domain, vinculin binds, and F-actin is recruited. The effect of actin is an increase in the force on talin by 3 pN, which is the tension measured on single vinculin-actin linkages [24]. Hence, vinculin binding and actin recruitment initially operate as a positive feedback, promoting talin unfolding and further vinculin binding. However, as the force across talin keeps increasing, vinculin molecules start to unbind, regulating the force level to a stable value around 23 pN. This force is determined by the equilibrium between vinculin binding and unbinding rates, and do not depend on the magnitude of the mechanical reinforcement by actin filaments. Interestingly, single-molecule assays have suggested that the integrin-fibronectin association operates as a catch bond, with an optimal lifetime at forces in the range of those predicted by our model [39].

Our simplified model overlooks the complexity of focal adhesions, which do not only involve talin-mediated linkages and include a multitude of interacting actors that operate in the different phases of its maturation and function [4, 21]. For instance, while all 13 talin rod domains can unfold under force, they have hierarchical stability, which suggests a range of mechanical thresholds for gradual vinculin recruitment [22]. Additionally, cytoplasmatic vinculin exists in an autoinhibited state, which would likely decrease its effective affinity for talin. Talin is a mechanosensing hub that recruits many other binding partners, which could alter its folding properties to regulate the mechanical response of the adhesion [11]. For example, RIAM and DLC1, unlike vinculin, bind folded talin bundles, which could stabilize talin and increase the force threshold for vinculin recruitment. Altogether, the integration of our data with these additional factors would extend the range of regulatory mechanisms in these mechanosensitive linkages, which could explain the broad range of integrin loads observed, and the variety of mechanical responses observed in cells [42].

In summary, the observation of the coil-to-helix contraction that occurs upon vinculin binding has allowed us to describe how the interaction between vinculin and talin is regulated by force, and which could be its implications for force transmission in focal adhesions. While previous work demonstrated that talin unfolding by force was necessary for force transmission and transduction in focal adhesions [43], how vinculin binding regulates this cellular process remained an open question. Vinculin transmits forces in focal adhesions [24], and there are at least 11 vinculin sites in each talin molecule. Hence, there is a clear force pathway for the increase in tension along talin, given by gradual vinculin recruitment. However, it remained unclear how this force level could be regulated, especially since vinculin is an indicator of stable and mature focal adhesions [17, 24]. Our results have demonstrated that the mechanics of the talin-vinculin interaction define a negative feedback by which the mechanical homeostasis of focal adhesions could be maintained to form stable cell adhesions. Hence, our findings provide a first basis by which the talin-vinculin interaction could work in cellular adhesions to regulate force transmission and transduction.

## IV. MATERIALS AND METHODS

### A. Magnetic Tweezers Setup

All experiments were done on our custom-made magnetic tweezers setup, as described before [26, 27]. Single molecules were tethered to superparamagnetic Dynabeads M-270 beads (2.8 *µ*m diameter). Calibrated forces were applied using either a voice-coil mounted pair of permanent magnets (Equipment Solutions), or a magnetic tape head (BRUSH 902836). Image processing was done by custom-written software written in C++/Qt, fully available on the lab’s website (http://zeptowatt.com). All experiments are done in custom-made fluid chambers built by two sandwiched glass coverslips, separated by a laser-cut parafilm patter. The fluid chambers are functionalized to add the HaloTag ligand and reference beads as described before [26]. Both the fluid chambers and magnetic beads are passivized using TRIS blocking buffer (20 mM Tris-HCL pH 7.4, 150 mM NaCl, 2mM MgCl2, and 1% w/v/ sulfhydryl blocked-BSA). All experiments are carried out in HEPES buffer (Hepes 10 mM pH 7.2, NaCl 150 mM, EDTA 1 mM), and the desired vinculin head concentration. See SI (Section XIII) for further details.

### B. Protein expression and purification

Polyprotein constructs are engineered using of BamHI, BglII, and KpnI restriction sites in pFN18a restriction vector, as described previously [26]. Our protein construct contains the R3 IVVI, or R3 WT mouse talin domain, followed by eight titin I91 domains, and flanked by an N-terminal HaloTag enzyme and a C-terminal AviTag for biotinylation. Human vinculin head was expressed and purified following an analogous procedure, skipping the biotinylation process. See SI (Section XIII) for further details.

### C. Single-molecule data analysis

Our data acquisition software collects data as a binary file, visualized later with custom-written software in Igor Pro (Wavemetrics). All data is acquired at 1,000 to 1,600 frames per second. Data is smoothed with a 4th order Savitzky-Golay filter using a box size of N=101. The folding/unfolding states of talin are automatically detected using a double threshold algorithm. See SI (Section XIII) for further details.

## ACKNOWLEDGMENTS

We thank Dr. Igor Barsukov from University of Liverpool for sharing the R3 plasmid with us. We thank all of the members of the J.M.F. laboratory for the valuable discussions and comments on the manuscript.

## Author contributions

R.T-R., A. A-C., and J.M.F. designed research; R.T-R., and A.A-C. conducted experiments and analyzed the data; R.T-R. conducted the computer simulations; R.T-R. and J.M.F. wrote the paper.

## REFERENCES

[1] E. C. Yusko and C. L. Asbury, “Force is a signal that cells cannot ignore,” Molecular Biology of the Cell, vol. 25, no. 23, pp. 3717–3725, 2014.

[2] P. Roca-Cusachs, V. Conte, and X. Trepat, “Quantifying forces in cell biology,” Nature Cell Biology, vol. 19, pp. 742–751, 2017.

[3] J. Eyckmans, T. Boudou, X. Yu, and C. S. Chen, “A hitchhiker’s guide to mechanobiology,” Developmental Cell, vol. 21, no. 1, pp. 35–47, 2011.

[4] A. Elosegui-Artola, X. Trepat, and P. Roca-Cusachs, “Control of mechanotransduction by molecular clutch dynamics,” Trends in Cell Biology, vol. 28, no. 5, pp. 356–367, 2018.

[5] K. Burridge and M. Chrzanowska-Wodnicka, “Focal adhesions, contractility, and signaling,” Annual Review of Cell and Developmental Biology, vol. 12, no. 1, pp. 463–519, 1996.

[6] V. P. Hytönen and B. Wehrle-Haller, “Mechanosensing in cell–matrix adhesions – converting tension into chemical signals,” Experimental Cell Research, vol. 343, no. 1, pp. 35–41, 2016.

[7] K. R. Legate and R. Fassler, “Mechanisms that regulate adaptor binding to *β*-integrin cytoplasmic tails,” Journal of Cell Science, vol. 122, no. 2, pp. 187–198, 2009.

[8] K. Burridge and L. Connell, “A new protein of adhesion plaques and ruffling membranes,” The Journal of Cell Biology, vol. 97, no. 2, pp. 359–367, 1983.

[9] S. J. Shattil, C. Kim, and M. H. Ginsberg, “The final steps of integrin activation: the end game,” Nature Reviews Molecular Cell Biology, vol. 11, pp. 288–300, 2010.

[10] P. Kanchanawong, G. Shtengel, A. M. Pasapera, E. B. Ramko, M. W. Davidson, H. F. Hess, and C. M. Waterman, “Nanoscale architecture of integrin-based cell adhesions,” Nature, vol. 468, pp. 580–584, 2010.

[11] B. T. Goult, T. Zacharchenko, N. Bate, R. Tsang, F. Hey, A. R. Gingras, P. R. Elliott, G. C. K. Roberts, C. Ballestrem, A. R. Critchley, and I. L. Barsukov, “Riam and vinculin binding to talin are mutually exclusive and regulate adhesion assembly and turnover,” Journal of Biological Chemistry, vol. 288, no. 12, pp. 8238–8249, 2013.

[12] G. Jiang, G. Giannone, D. R. Critchley, E. Fukumoto, and M. P. Sheetz, “Two-piconewton slip bond between fibronectin and the cytoskeleton depends on talin,” Nature, vol. 424, no. 6946, pp. 334–337, 2003.

[13] R. Zaidel-Bar, S. Itzkovitz, A. Ma’ayan, R. Iyengar, and B. Geiger, “Functional atlas of the integrin adhesome,” Nature Cell Biology, vol. 9, pp. 858–867, 2007.

[14] B. T. Goult, J. Yan, and M. A. Schwartz, “Talin as a mechanosensitive signaling hub,” The Journal of Cell Biology, vol. 217, no. 11, pp. 3776–3784, 2018.

[15] A. R. Gingras, W. H. Ziegler, R. Frank, I. L. Barsukov, G. C. K. Roberts, D. R. Critchley, and J. Emsley, “Mapping and consensus sequence identification for multiple vinculin binding sites within the talin rod,” Journal of Biological Chemistry, vol. 280, no. 44, pp. 37217–37224, 2005.

[16] A. del Rio, R. Perez-Jimenez, R. Liu, P. Roca-Cusachs, J. M. Fernandez, and M. P. Sheetz, “Stretching single talin rod molecules activates vinculin binding,” Science, vol. 323, no. 5914, pp. 638–641, 2009.

[17] C. Ciobanasu, B. Faivre, and C. Le Clainche, “Actomyosin-dependent formation of the mechanosensitive talin–vinculin complex reinforces actin anchoring,” Nature Communications, vol. 5, p. 3095, 2014.

[18] A. Carisey, R. Tsang, A. M. Greiner, N. Nijenhuis, N. Heath, A. Nazgiewicz, R. Kemkemer, B. Derby, J. Spatz, and C. Ballestrem, “Vinculin regulates the recruitment and release of core focal adhesion proteins in a force-dependent manner,” Current Biology, vol. 23, no. 4, pp. 271–281, 2013.

[19] J. D. Humphries, P. Wang, C. Streuli, B. Geiger, M. J. Humphries, and C. Ballestrem, “Vinculin controls focal adhesion formation by direct interactions with talin and actin,” The Journal of Cell Biology, vol. 179, no. 5, pp. 1043–1057, 2007.

[20] H. Hirata, H. Tatsumi, C. T. Lim, and M. Sokabe, “Force-dependent vinculin binding to talin in live cells: a crucial step in anchoring the actin cytoskeleton to focal adhesions,” American Journal of Physiology-Cell Physiology, vol. 306, no. 6, pp. C607–C620, 2014.

[21] L. B. Case and C. M. Waterman, “Integration of actin dynamics and cell adhesion by a three-dimensional, mechanosensitive molecular clutch,” Nature Cell Biology, vol. 17, pp. 955–963, 2015.

[22] A. W. M. Haining, M. von Essen, S. J. Attwood, V. P. Hytönen, and A. del Río Hernández, “All subdomains of the talin rod are mechanically vulnerable and may contribute to cellular mechanosensing,” ACS Nano, vol. 10, pp. 6648–6658, 07 2016.

[23] M. Yao, B. T. Goult, B. Klapholz, X. Hu, C. P. Toseland, Y. Guo, P. Cong, M. P. Sheetz, and J. Yan, “The mechanical response of talin,” Nature Communications, vol. 7, p. 11966, 2016.

[24] C. Grashoff, B. D. Hoffman, M. D. Brenner, R. Zhou, M. Parsons, M. T. Yang, M. A. McLean, S. G. Sligar, C. S. Chen, T. Ha, and M. A. Schwartz, “Measuring mechanical tension across vinculin reveals regulation of focal adhesion dynamics,” Nature, vol. 466, pp. 263–266, 2010.

[25] M. Yao, B. T. Goult, H. Chen, P. Cong, M. P. Sheetz, and J. Yan, “Mechanical activation of vinculin binding to talin locks talin in an unfolded conformation,” Scientific Reports, vol. 4, p. 4610, 2014.

[26] I. Popa, J. A. Rivas-Pardo, E. C. Eckels, D. J. Echelman, C. L. Badilla, J. Valle-Orero, and J. M. Fernández, “A halotag anchored ruler for week-long studies of protein dynamics,” Journal of the American Chemical Society, vol. 138, pp. 10546–10553, 08 2016.

[27] R. Tapia-Rojo, E. C. Eckels, and J. M. Fernández, “Ephemeral states in protein folding under force captured with a magnetic tweezers design,” Proceedings of the National Academy of Sciences, vol. 116, no. 16, pp. 7873–7878, 2019.

[28] E. Papagrigoriou, A. R. Gingras, I. L. Barsukov, N. Bate, I. J. Fillingham, B. Patel, R. Frank, W. H. Ziegler, G. C. K. Roberts, D. R. Critchley, and J. Emsley, “Activation of a vinculin-binding site in the talin rod involves rearrangement of a five-helix bundle,” The EMBO journal, vol. 23, pp. 2942–2951, 08 2004.

[29] T. Izard, G. Evans, R. A. Borgon, C. L. Rush, G. Bricogne, and P. R. J. Bois, “Vinculin activation by talin through helical bundle conversion,” Nature, vol. 427, no. 6970, pp. 171–175, 2004.

[30] M. Doi and S. Edwards, The Theory of Polymer Dynamics. Oxford Science Publications, 1986.

[31] B. P. English, W. Min, A. M. van Oijen, K. T. Lee, G. Luo, H. Sun, B. J. Cherayil, S. C. Kou, and X. S. Xie, “Ever-fluctuating single enzyme molecules: Michaelis-menten equation revisited,” Nature Chemical Biology, vol. 2, no. 2, pp. 87–94, 2006.

[32] S. C. Kou, B. J. Cherayil, W. Min, B. P. English, and X. S. Xie, “Single-molecule michaelis menten equations,” The Journal of Physical Chemistry B, vol. 109, no. 41, pp. 19068–19081, 2005.

[33] M. D. Bass, B. Patel, I. G. Barsukov, I. J. Fillimgham, R. Mason, B. J. Smith, C. R. Bagshaw, and D. R. Critchley, “Further characterization of the interaction between the cytoskeletal proteins talin and vinculin,” Biochemical Journal, vol. 362, no. 3, pp. 761–768, 2002.

[34] G. Bell, “Models for the specific adhesion of cells to cells,” Science, vol. 200, no. 4342, pp. 618–627, 1978.

[35] R. Merkel, P. Nassoy, A. Leung, K. Ritchie, and E. Evans, “Energy landscapes of receptor–ligand bonds explored with dynamic force spectroscopy,” Nature, vol. 397, no. 6714, pp. 50–53, 1999.

[36] M. Rief, M. Gautel, F. Oesterhelt, J. M. Fernandez, and H. E. Gaub, “Reversible unfolding of individual titin immunoglobulin domains by afm,” Science, vol. 276, no. 5315, pp. 1109–1112, 1997.

[37] A. P. Wiita, S. R. K. Ainavarapu, H. H. Huang, and J. M. Fernandez, “Force-dependent chemical kinetics of disulfide bond reduction observed with single-molecule techniques,” Proceedings of the National Academy of Sciences, vol. 103, no. 19, pp. 7222–7227, 2006.

[38] W. Thomas, M. Forero, O. Yakovenko, L. Nilsson, P. Vicini, E. Sokurenko, and V. Vogel, “Catch-bond model derived from allostery explains force-activated bacterial adhesion,” Biophysical Journal, vol. 90, no. 3, pp. 753–764, 2006.

[39] F. Kong, A. J. García, A. P. Mould, M. J. Humphries, and C. Zhu, “Demonstration of catch bonds between an integrin and its ligand,” The Journal of Cell Biology, vol. 185, no. 7, pp. 1275–1284, 2009.

[40] I. Heller, T. P. Hoekstra, G. A. King, E. J. G. Peterman, and G. J. L. Wuite, “Optical tweezers analysis of dna–protein complexes,” Chemical Reviews, vol. 114, pp. 3087–3119, 03 2014.

[41] M. Huse, “Mechanical forces in the immune system,” Nature Reviews Immunology, vol. 17, pp. 679–690, 07 2017.

[42] H. P. Erickson, “Protein unfolding under isometric tension—what force can integrins generate, and can it unfold fniii domains?,” Current Opinion in Structural Biology, vol. 42, pp. 98–105, 2017. Folding and binding * Proteins: Bridging theory and experiment.

[43] A. Elosegui-Artola, R. Oria, Y. Chen, A. Kosmalska, C. Pérez-González, N. Castro, C. Zhu, X. Trepat, and P. Roca-Cusachs, “Mechanical regulation of a molecular clutch defines force transmission and transduction in response to matrix rigidity,” Nature Cell Biology, vol. 18, pp. 540–548, 2016.

