## Supplementary Information for "Direct Observation of a Coil-to-Helix Contraction Triggered by Vinculin Binding to Talin"

Rafael Tapia-Rojo,[1] Alvaro Alonso-Caballero,[1] and Julio M. Fernandez[1]

[1]*Department of Biological Sciences, Columbia University, New York, NY 10027, USA*

### I. STRUCTURAL MODEL OF THE TALIN ROD DOMAINS

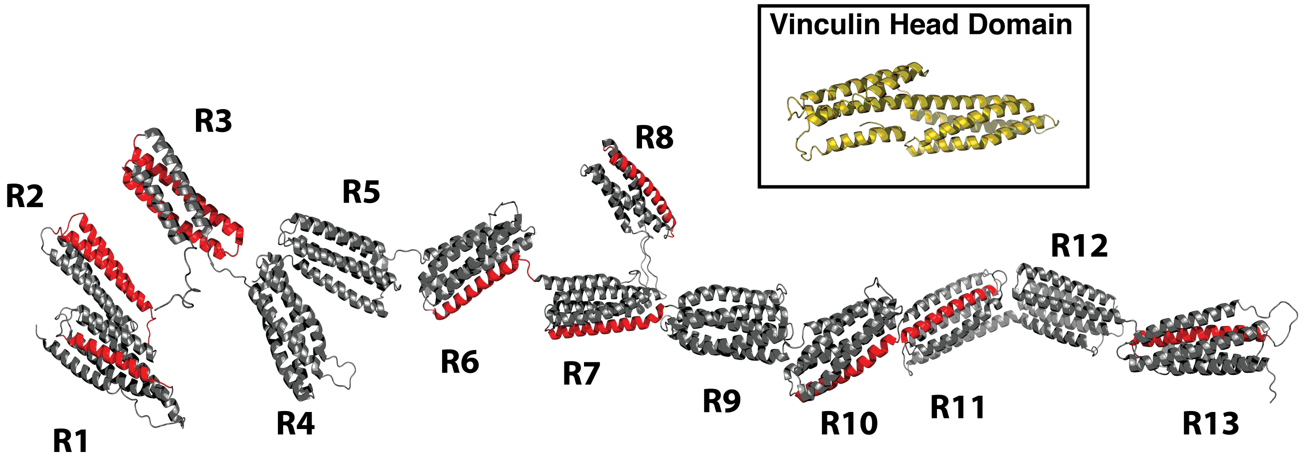

FIG. 1. **Talin rod domains and Vinculin Head:** The mechanosensing region of talin has 13 four or five $\alpha$-helix bundle domains, with at least 11 vinculin binding sites (red helices). The unfolding of talin domains exposes these cryptic sites, which activates vinculin by displacing the interaction between its head and tail, binding to the helices with high affinity. (Inset) Vinculin head.

### II. MECHANICAL CHARACTERIZATION OF TALIN R3 IVVI DOMAIN

We characterize the mechanics of the R3 WT and R3 IVVI domains through folded/unfolded extension change (step-sizes), the folding probability, and the folding/unfolding rates (Fig. S2). The step-sizes are equal for both domains, Fig. S2A. They scale with force following the Freely-Jointed Chain (FJC) model of polymer elasticity, with a contour length increase of $L_c = 39.81 \pm 9.73$ nm, and a Kuhn length of $l_K = 0.77 \pm 0.27$ nm. The R3 domain has 124 residues, so the value reported is reasonable. The folding probability is the relative occupation of the folded state (Fig. S2B), and follows a sigmoidal function $P_f(F) = \left(\exp\left(\frac{F-F_{1/2}}{r}\right)+1\right)^{-1}$. The mechanical stabilities of R3 WT and R3 IVV are very different; R3 WT has a coexistence force of $F_{1/2} = 5.14 \pm 0.08$ pN, while R3 IVVI $F_{1/2} = 9.09 \pm 0.02$ pN. The folding and unfolding rates are calculated from the inverse of the average dwell times, as determined from exponential fits to the dwell time distributions (Fig. S2C). Due to the small force range over which R3 operates, they follow a simple Bell-Evans model. The R3 WT and R3 IVVI have very different kinetics, yielding: $x^\dagger_F = 6.42 \pm 0.28$ nm, $k^F_0 = (1.84 \pm 1.01) \times 10^6$ s$^{-1}$, and $x^\dagger_U = 5.54 \pm 0.28$ nm, and $k^U_0 = (4.61 \pm 2.52) \times 10^{-6}$ s$^{-1}$ (R3 IVVI); and $k^F_0 = (1.2 \pm 0.8) \times 10^5$ s$^{-1}$, and $x^\dagger_U = 4.85 \pm 0.56$ nm, and $k^U_0 = (2.50 \pm 1.01) \times 10^{-3}$ s$^{-1}$ (R3 WT).

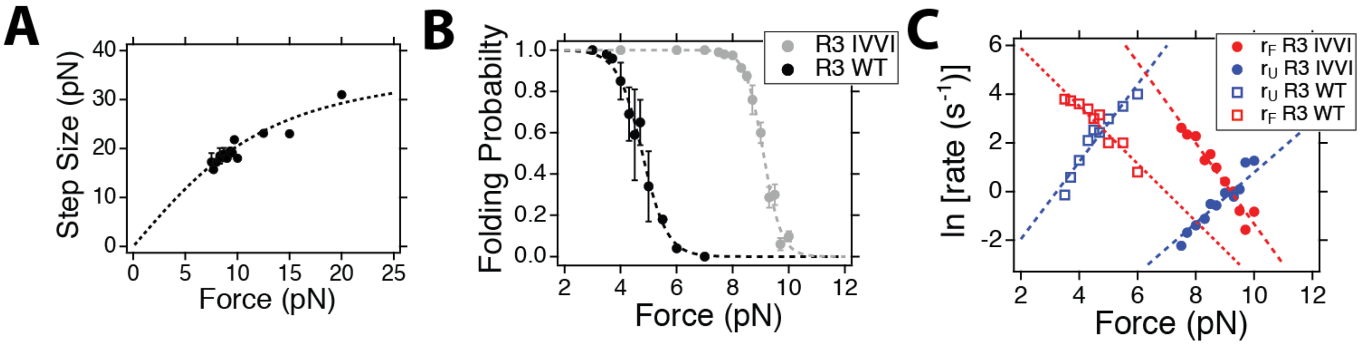

FIG. 2. **Mechanical properties of talin R3 domain:** (A) Step sizes as a function of the force, fitted to a FJC model (dotted line). (B) Folding probability fitted to a sigmoidal function (dotted line). (C) Folding (red) and unfolding (blue) rates, fitted to a Bell model.

### III. HOURS-LONG RECORDINGS IN THE ABSENCE AND PRESENCE OF VINCULIN HEAD

Magnetic tweezers have high stability, which allows measuring the same molecule over several hours. To further clarify the fingerprint for vinculin head binding, we measure the dynamics of the R3 IVVI domain in the absence of vinculin head, and with 30 nM vinculin head (Fig. S3). In the absence of vinculin head, at a force of 9 pN, the talin R3 IVVI domain hops continually between its folded and unfolded state; over more than 5 hours, the dynamics of talin remain unaltered. In the presence of 30 nM vinculin head, binding occurs in a few seconds, and talin is locked on its unfolded state for the rest of the experiment. We have used long recordings in the absence of vinculin (5, 7, and 36 hours) to estimate an upper bound for the rate of binding in absence of vinculin head (1.7 $\times 10^{-5}$ s$^{-}$1).

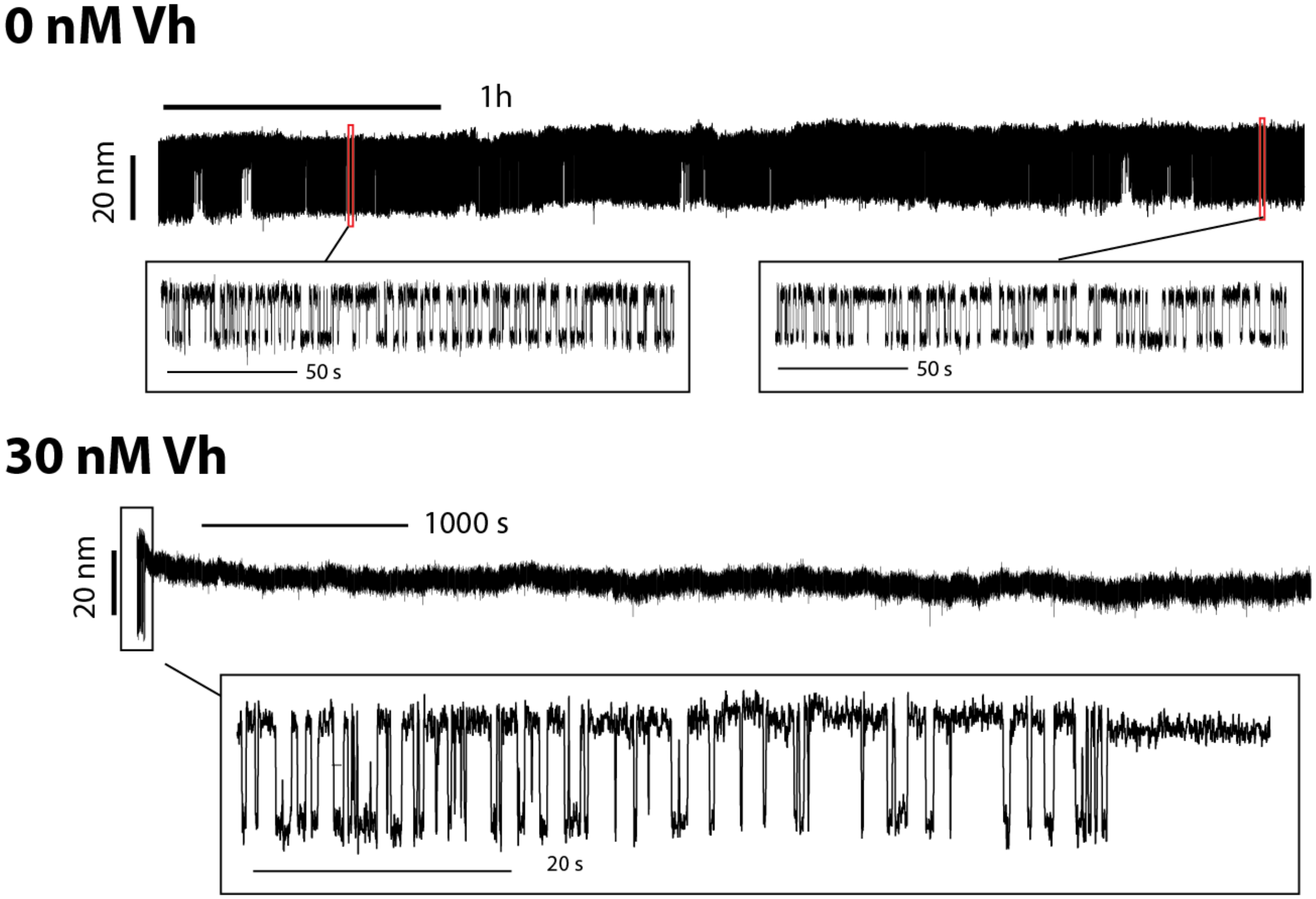

FIG. 3. **Hours-long recordings of R3 IVVI in the absence and presence of vinculin head:** R3 IVVI maintains its folding dynamics unaltered for several hours (upper). In the presence of vinculin head, these dynamics cease after a few seconds, and the R3 domain remains locked on its unfolded conformation.

### IV. THE VINCULIN BINDING FINGERPRINT

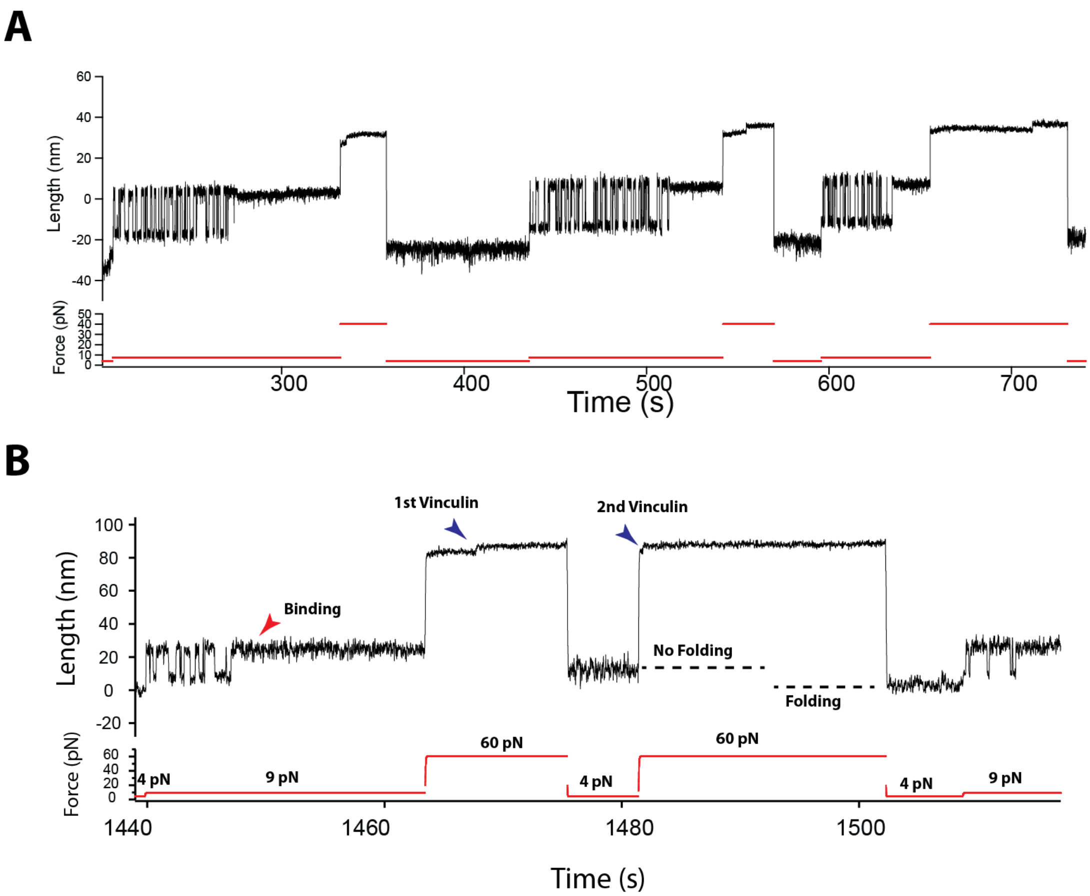

FIG. 4. **The vinculin binding fingerprint:** (A) Typical vinculin binding experiment, consisting of a repetition of three folding pulses. The cycle starts at 4 pN (reference force), where R3 IVVI is folded. The probe pulse (9 pN in this case) is used to detect vinculin binding in the arrest of talin folding dynamics. Finally, vinculin head is expelled through a high force pulse (> 40 pN). This protocol is cycled. (B) Example trajectory with two separated unbinding events. We show here that the two vinculin molecules must unbind for talin to recover the unbound state, characterized by talin ability to refold under force.

### V. VINCULIN HEAD BINDING TO THE R3 WT DOMAIN

The R3 WT domain has a lower mechanical stability than the mutant R3 IVVI due to the presence of a four-threonine belt in its hydrophobic core that, however, do not affect its vinculin-binding properties. We have investigated vinculin head binding to R3 WT to demonstrate the generality of our measurements and the mechanism for force regulation we propose. Figure S5 summarizes our observations. The fingerprint for vinculin head binding is the same for R3 WT and R3 IVVI; R3 folding dynamics are arrested due to vinculin head binding, which is characterized by a contraction of the R3 polypeptide due to a coil-to-helix transition. Figures S5A and B show two binding recordings at forces of 5 and 12.5 pN. The coexistence force for R3 WT is 5 pN, compared to 9 pN for R3 IVVI, which impedes to resolve the binding contraction; however, it is clearly observed at higher forces like 12.5 pN. We measure the binding contraction and unbinding steps, which follow the same force dependency we obtained for R3 IVVI (Fig. S5C). This indicates that R3 WT experiences the same coil-to-helix transition due to the binding of two vinculin head molecules. We also measured the binding probability, which shows the same shape we measured for R3 IVVI, and can be described with our theoretical model. The threshold for binding decreases to 4 pN due to the lower mechanical stability, but the negative force dependence is equivalent to that of R3 IVVI, since the contraction steps are equal.

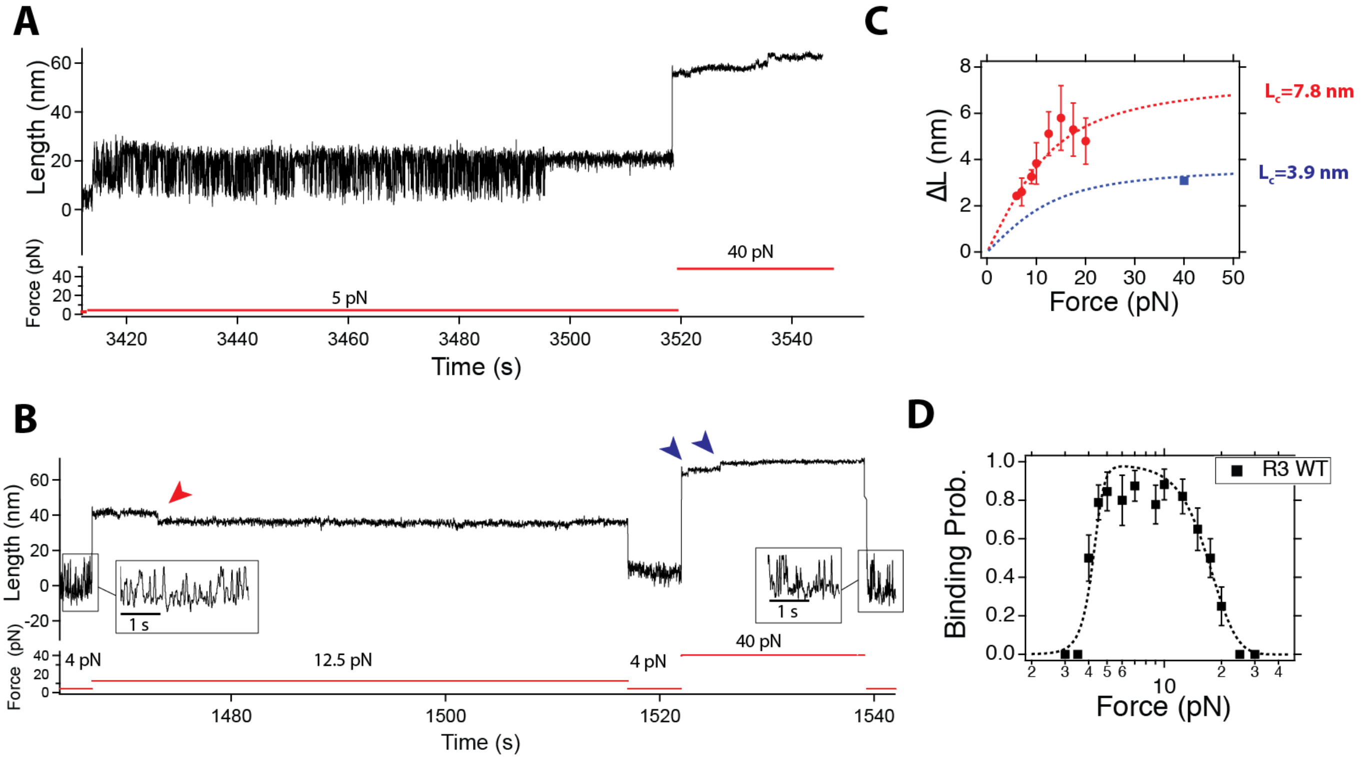

FIG. 5. **Vinculin head binding to R3 WT:** (A, B) Magnetic tweezers recordings of the talin R3 WT domain at 20 nM vinculin head. The fingerprint for vinculin head binding is the same as with R3 IVVI, only that binding occurs at lower forces due to the lower mechanical stability of R3 WT. (C) Force-dependence of the binding contraction (red) and unbinding steps (blue), which follow the same FJC measured for R3 IVVI. This indicates that vinculin head binding triggers the same coil-to-helix transition on R3 WT. (D) Binding probability of vinculin head to the R3 WT domain.

### VI. UNBINDING KINETICS

Vinculin unbinding is triggered by forces above 40 pN. This results in a sequential dissociation of the two bound vinculin heads through ∼3 nm steps, due to the uncoiling of each of the two vinculin-binding site helices. The kinetics of unbinding depend steeply on the pulling force; at 40 pN several hundreds of seconds are needed to liberate talin, while at 60 pN unbinding takes place in a few seconds. In the majority of our recordings, we observe two distinct steps that indicate the sequential unbinding. In a fraction of trajectories (40%), only one step is observed. This is likely due to very fast unbinding events that are buried in the elastic equilibration or occur too fast to be resolved. Figure S6A shows an unbinding recording at 40 pN, where the first step occurs within 1 second, while the second one needs over 200 seconds.

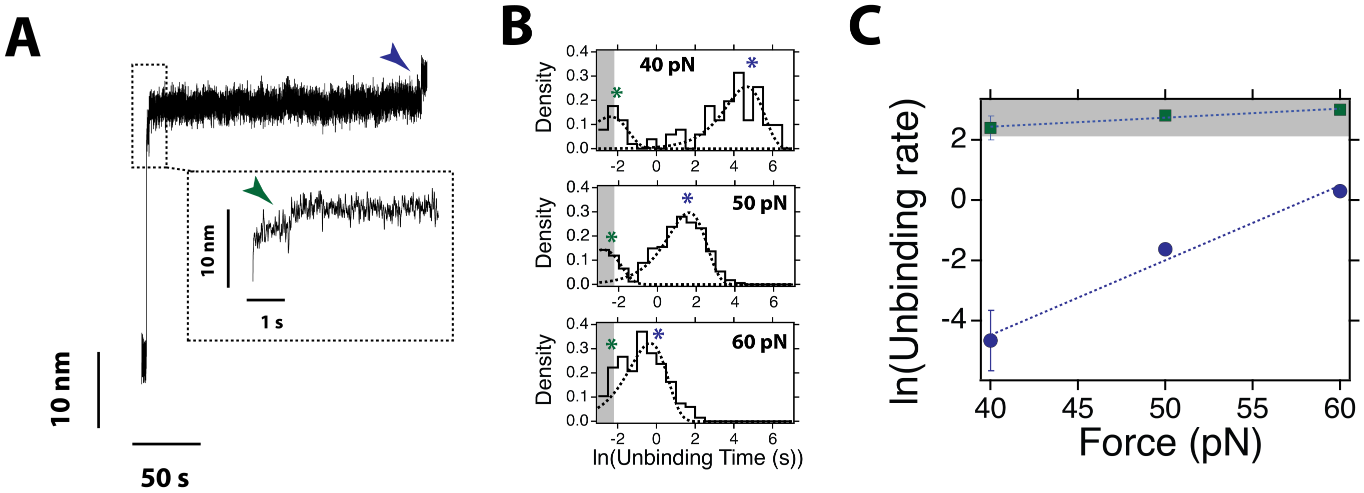

FIG. 6. **Vinculin unbinding kinetics**: (A) Typical unbinding trajectory at a force of 40 pN. The two unbinding events can occur in different kinetic timescales. We observe a first one occurring within the first second (green arrow), and a second much slower one occurring after hundreds of seconds. (B) Logarithmic dwell time histograms for the unbinding kinetics at 40, 50, and 60 pN. The logarithmic scale allows separating different concurrent kinetic processes since exponential distributions are transformed into peaked distributions. At the three forces, a bimodal distribution is obtained, with fast events within a few milliseconds (green stars) and slow events occurring in seconds (blue). The fast events occur within our limit of detection upon force changes (grey bar, ∼100 ms). (C) Force dependency of the unbinding rates for the fast (green) and slow (blue) events, as obtained from the dwell-time distributions. The force dependency of the fast events could not be accurately determined since it enters on the limit of detection, set at 100 ms. The fast events show a steep force dependency, following Bell model with a distance to the transition state of $x^{\dagger} = 1.05$ nm.

We measure the unbinding kinetics at 40, 50, and 60 pN. Figure S6B shows the distributions of unbinding times calculated with logarithmic binning. The logarithmic histograms suggest two separate unbinding kinetics, with a fast one occurring within the detection level (green star), and a slow one that depends strongly on the pulling force (blue star). The histograms are fitted to a double exponential distribution to extract the average unbinding times. Respectively, they were constructed with 101 (40 pN), 335 (50 pN), and 134 (60 pN) unbinding events. We plot the unbinding rate as a function of the force in Fig. S6C, as obtained from the dwell-time histogram fits. The fast kinetics (green) enter on the detection limit (grey area), so their force dependency cannot be fully resolved. The slow unbinding kinetics depend on the force following a typical Bell exponential model, yielding a distance to transition state of $x^{\dagger} = 1.05$ nm, and a prefactor $k_{\mathrm{ub}} = 5.6 \times 10^{-7}$ s$^{-1}$.

### VII. KINETICS OF THE BINDING CONTRACTION

The contraction of the R3 polypeptide that arises from vinculin binding is not an instantaneous event, but a slow relaxation that accelerates with force. At low forces, it occurs within hundreds of milliseconds, while above 15 pN it is a fast transition. Figure S7A shows averaged contractions fitted to an exponential decay function, which allows estimating the timescale at which the binding event occurs. Figure S7B shows the dependence of the binding contraction kinetics as a function of force. These kinetics suggest that vinculin head binding likely requires a maturation process, perhaps arising from sequential structural rearrangements required for vinculin recognition, accommodation, and binding. The acceleration of the binding kinetics with force is likely related to the evolution of the underlying free energy landscape of unfolded talin, which is dominated by polymer elasticity. The width of the free energy basin of unfolded talin decreases with force; hence, we could relate the acceleration of the contraction kinetics with the diffusion over the free energy barrier. However, the specifics of the actual landscape over which the coil-to-helix transition occurs are unknown.

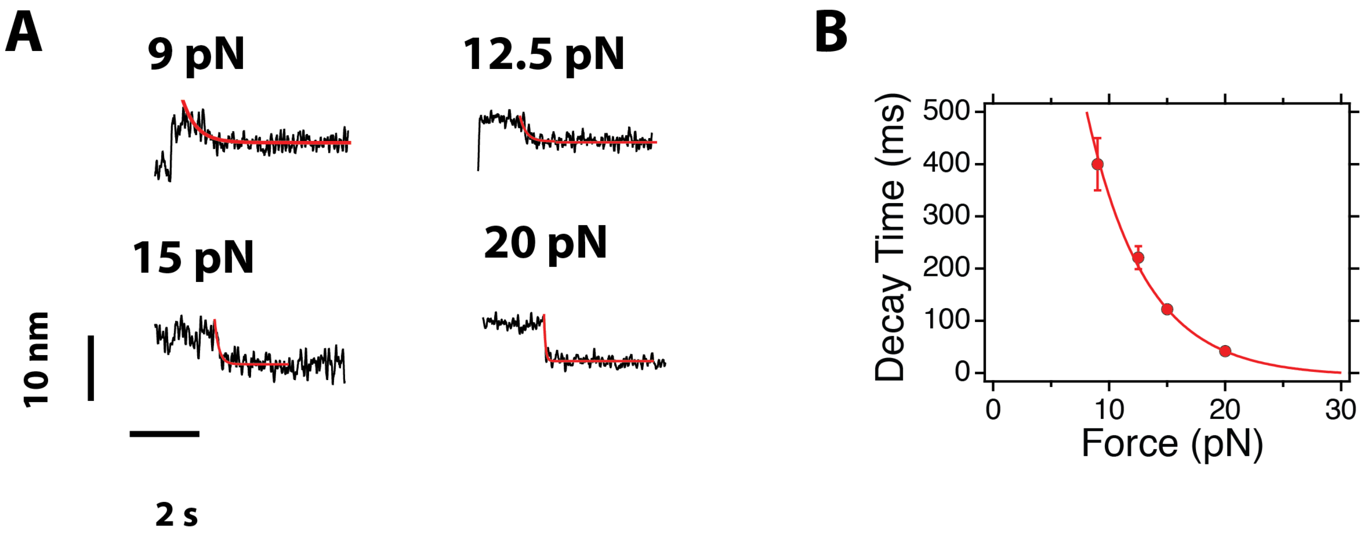

FIG. 7. **Kinetics of the vinculin binding event to R3 IVVI:** (A) Average contraction of the talin polypeptide upon vinculin head binding. The contraction is fitted to an exponential decay function, showing the acceleration of the kinetics with the force. (B) Time constant of the contraction event as a function of the force.

### VIII. VINCULIN HEAD CONCENTRATION

In our preparation of vinculin head (see Methods), we observe a strong tendency to aggregate. This might decrease the fraction of active vinculin head molecules that can bind, leading to discrepancies in the concentration we measure. Here, we have used three independent preparations of vinculin head, which show small differences in the binding rate at the same concentration. Figure S8 shows the binding rate of vinculin head for each of the three preparations, which, however, all show a quadratic dependence.

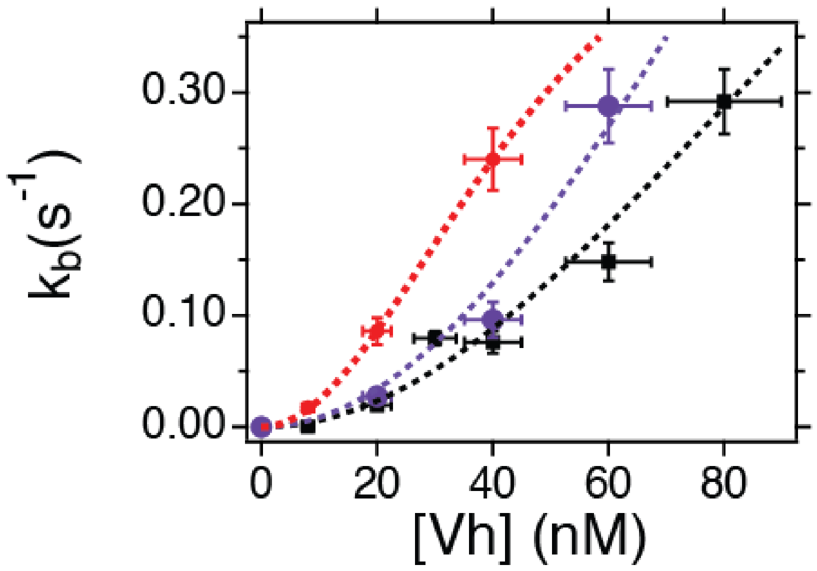

FIG. 8. **Binding rate of vinculin head for three different molecular preparations**

### IX. SINGLE MOLECULE BINDING KINETICS OF VINCULIN TO UNFOLDED TALIN

We model vinculin binding to unfolded talin with the following kinetic model:

$$\mathrm{F} \underset{\mathrm{k_F}}{\overset{\mathrm{k_U}}{\rightleftharpoons}} \mathrm{U} + 2\mathrm{Vh} \xrightarrow{k^0_{\mathrm{on}}} \mathrm{U_b},$$

Here, we assume that talin folds and unfolds in equilibrium with respective rates $k_{\mathrm{F}}$ and $k_{\mathrm{U}}$, that vinculin only binds to unfolded talin, and that two vinculin molecules are necessary to form the complex, so that the pseudo-binding constant can be written as $k^0_{\mathrm{on}} = k_{\mathrm{on}}[Vh]^2$, being $[Vh]$ the concentration of vinculin head. Here, we assume that the rates of unbinding are negligible, which is true over the force range where we observe binding. This model leads to the following set of linear differential equations:

$$\frac{\mathrm{d}P_{\mathrm{F}}}{\mathrm{d}t} = k_{\mathrm{F}}P_{\mathrm{U}} - k_{\mathrm{U}}P_{\mathrm{F}} \tag{1}$$

$$\frac{\mathrm{d}P_{\mathrm{U}}}{\mathrm{d}t} = k_{\mathrm{U}}P_{\mathrm{F}} - (k_{\mathrm{F}} + k^0_{\mathrm{on}})P_{\mathrm{U}} \tag{2}$$

$$\frac{\mathrm{d}P_{\mathrm{b}}}{\mathrm{d}t} = k^0_{\mathrm{on}}P_{\mathrm{U}} \tag{3}$$

Here, $P_{\mathrm{F}}$, $P_{\mathrm{U}}$, and $P_{\mathrm{b}}$ are the occupation probabilities of folded, unfolded talin, and bound state, respectively. The experimentally relevant observable is the waiting time to reach the bound state, $t_{\mathrm{b}}$. Its distribution $f(t_{\mathrm{b}})$ can be written as:

$$f(t_b) = \frac{\mathrm{d}P_{\mathrm{b}}}{\mathrm{d}t} = k^0_{\mathrm{on}}P_{\mathrm{U}} = k_{\mathrm{on}}[V]^2 P_{\mathrm{U}}. \tag{4}$$

Solving the kinetic model, we obtain an analytical expression for $f(t_{\mathrm{b}})$ assuming that our initial conditions are $P_{\mathrm{F}}(0) = 1$ (we start with folded talin), $P_{\mathrm{U}}(0) = 0$, $P_{\mathrm{b}}(0) = 0$, and the constraint $P_{\mathrm{F}}(t) + P_{\mathrm{U}}(t) + P_{\mathrm{b}}(t) = 1$:

$$f(t_b) = \frac{k^0_{\mathrm{on}}k_{\mathrm{U}}}{2\alpha}\left[e^{(\alpha+\beta)t} - e^{(\beta-\alpha)t}\right], \tag{5}$$

where:

$$\alpha = \sqrt{(k_{\mathrm{U}} + k_{\mathrm{F}} + k^0_{\mathrm{on}})^2/4 - k^0_b r_{\mathrm{U}}}$$

$$\beta = -(k_{\mathrm{U}} + k_{\mathrm{F}} + k^0_{\mathrm{on}})/2.$$

This expression is used in the manuscript to fit the experimental distributions of waiting times at different vinculin concentrations (Fig. 4B main manuscript).

From this expression, we calculate the rate of binding $k_{\mathrm{b}}$, which is the inverse of the first moment of the distribution $\langle t_{\mathrm{b}}\rangle = \int_0^\infty t f(t)\mathrm{d}t$, so that the observed rate of binding $k_{\mathrm{b}} = 1/\langle t_{\mathrm{b}}\rangle$ is:

$$k_{\mathrm{b}} = -\frac{(\alpha^2 - \beta^2)^2}{2\beta k^0_{\mathrm{on}}k_{\mathrm{U}}},$$

which can be rewritten as:

$$k_b = \frac{k_{\mathrm{U}}[V]^2}{[V]^2 + K_{\mathrm{M}}}, \tag{6}$$

where $K_{\mathrm{M}} = (r_{\mathrm{U}} + r_{\mathrm{F}})/k_{\mathrm{on}}$. This equation is the single-molecule analog for a second-order Hill model, where the role of the maximum velocity is taken by the unfolding rate of talin, which is the limiting rate of the system since it activates the substrate.

### X. FORCE DEPENDENCY

Binding of vinculin to talin is a force-dependent reaction as all the kinetic rates involved in the process are force-dependent. First, vinculin only binds to unfolded talin, and talin folding dynamics depend on the force exponentially. Second, the on-rate $k_{\text{on}}$ also depends on the force, since we observe slower binding kinetics as we increase the force in the range where talin is unfolded (10-25 pN).

As measured before, the folding and unfolding rates of talin can be modeled with a simple Bell-Evans exponential dependency (Fig. S2).

$$k_{\text{F}}(F) = k_0^F e^{-Fx_{\text{F}}^{\dagger}/kT} \tag{7}$$

$$k_{\text{U}}(F) = k_0^U e^{Fx_{\text{U}}^{\dagger}/kT}. \tag{8}$$

$$\tag{9}$$

The force dependence of $k_{\text{on}}$ can be modeled from the mechanical fingerprint of binding: the shortening of talin polypeptide due to the formation of the $\alpha$-helices upon vinculin binding. Contracting a stretched polymer requires an energetic cost since binding does mechanical work against the pulling force to shorten the polymer to a lower extension than that dictated by the FJC model. In this regard, we assume that the on-rate can be written as an Arrhenius-like rate:

$$k_{\text{on}}^0(F) = Ae^{-\Delta W(F)/kT}, \tag{10}$$

where $\Delta W(F)$ is the mechanical work needed to contract unfolded talin as a function of the force, and $A$ a prefactor, which depends on vinculin concentration $A = [V]^2 A'$. To a first approximation, we assume that the mechanical work is simply $\Delta W \approx F \cdot \Delta L$, where $\Delta L$ is the force-dependent contraction we measure experimentally, which has the shape of the FJC model. More complex expressions for the mechanical work can be written, specifically by accounting for the shape of the free-energy landscape of the FJC, and integrating the work done on a diffusing particle over such landscape to go from its equilibrium distance to the force-dependent contraction measured experimentally. However, no differences have been appreciated between the simple model, and its more complex forms.

Experimentally, we measure the binding probability over a time-window of 50 s as a function of the force. From our kinetic model, we can write an expression for this:

$$P_{\text{b}}(F;t) = 1 - \exp\left[-\frac{k_{\text{U}}(F)k_{\text{on}}(F)t}{k_{\text{U}}(F) + k_{\text{F}}(F) + k_{\text{on}}(F)}\right], \tag{11}$$

where the explicit dependence of each of the three rates involved has been discussed above. This is the expression we employ to describe the data in Fig. 5 (B) of the main manuscript.

### XI. PARAMETERS FOR THE FIT IN FIGURE 5 (BINDING PROBABILITY)

TABLE I. Parameters employed to describe the data for the R3 IVVI and R3 WT domain in Fig. 5B (main manuscript), and its origin.

| Parameter | Meaning | Value | Origin |
|---|---|---|---|
| $k_{\mathrm{U}}^{0}$ | Unfolding rate prefactor (Bell model) | $1.84 \times 10^{-6}$ s$^{-1}$ | Fixed (obtained from Fig. S2C) |
| $x_{\mathrm{U}}^{\dagger}$ | Distance to transition state from folded | 5.54 nm | Fixed (obtained from Fig. S2C) |
| $k_{\mathrm{F}}^{0}$ | Folding rate prefactor (Bell model) | $6.42 \times 10^{6}$ s$^{-1}$ | Fixed (obtained from Fig. S2C) |
| $x_{\mathrm{F}}^{\dagger}$ | Distance to transition state from unfolded | 6.42 nm | Fixed (obtained from Fig. S2C) |
| $[V]$ | Concentration of vinculin | 20 nM | Experimental condition |
| $t$ | Measuring time window | 50 s | Experimental condition |
| $\Delta L_{\mathrm{c}}$ | Contour length change for binding | 7.5 nm | Fixed (obtained from FJC fit) |
| $l_{\mathrm{K}}$ | Kuhn length for binding | $0.12 \pm 0.01$ nm | Free parameter |
| $A'$ | Prefactor for binding rate | $0.0011 \pm 0.0003$ s$^{-1}$ nM$^{-2}$ | Free parameter |
| $k_{\mathrm{U}}^{0}$ | Unfolding rate prefactor (Bell model) | 0.006 s$^{-1}$ | Fixed (obtained from Fig. S2C) |
| $x_{\mathrm{U}}^{\dagger}$ | Distance to transition state from folded | 6.52 nm | Fixed (obtained from Fig. S2C) |
| $k_{\mathrm{F}}^{0}$ | Folding rate prefactor (Bell model) | 3844.5 s$^{-1}$ | Fixed (obtained from Fig. S2C) |
| $x_{\mathrm{F}}^{\dagger}$ | Distance to transition state from unfolded | 4.87nm | Fixed (obtained from Fig. S2C) |
| $[V]$ | Concentration of vinculin | 20 nM | Experimental condition |
| $t$ | Measuring time window | 50 s | Experimental condition |
| $\Delta L_{\mathrm{c}}$ | Contour length change for binding | 7.8 nm | Fixed (obtained from FJC fit) |
| $l_{\mathrm{K}}$ | Kuhn length for binding | $0.05 \pm 3.07$ nm | Free parameter |
| $A'$ | Prefactor for binding rate | $0.00025 \pm 0.0006$ s$^{-1}$ nM$^{-2}$ | Free parameter |

### XII. COMPUTATIONAL MODEL FOR THE NEGATIVE FEEDBACK MECHANISM DEFINED BY TALIN-VINCULIN INTERACTION

We integrate our findings on a minimalistic model for the mechanical linkage between integrin and F-actin, which accounts for the talin-vinculin-actin linkage. We represent this model as a control block system in Fig. 9A. Force flows by acting positively on talin domains—unfolding them—but negatively on vinculin binding, since the reaction is hampered by force. If binding occurs, vinculin recruits F-actin filaments, which increase force transmission. We model actin effect on the linkage as a net gain on the force across talin. The system is governed by the folding/unfolding rates of talin, and the binding/unbinding rates of vinculin, which we have measured experimentally (Fig. S9B).

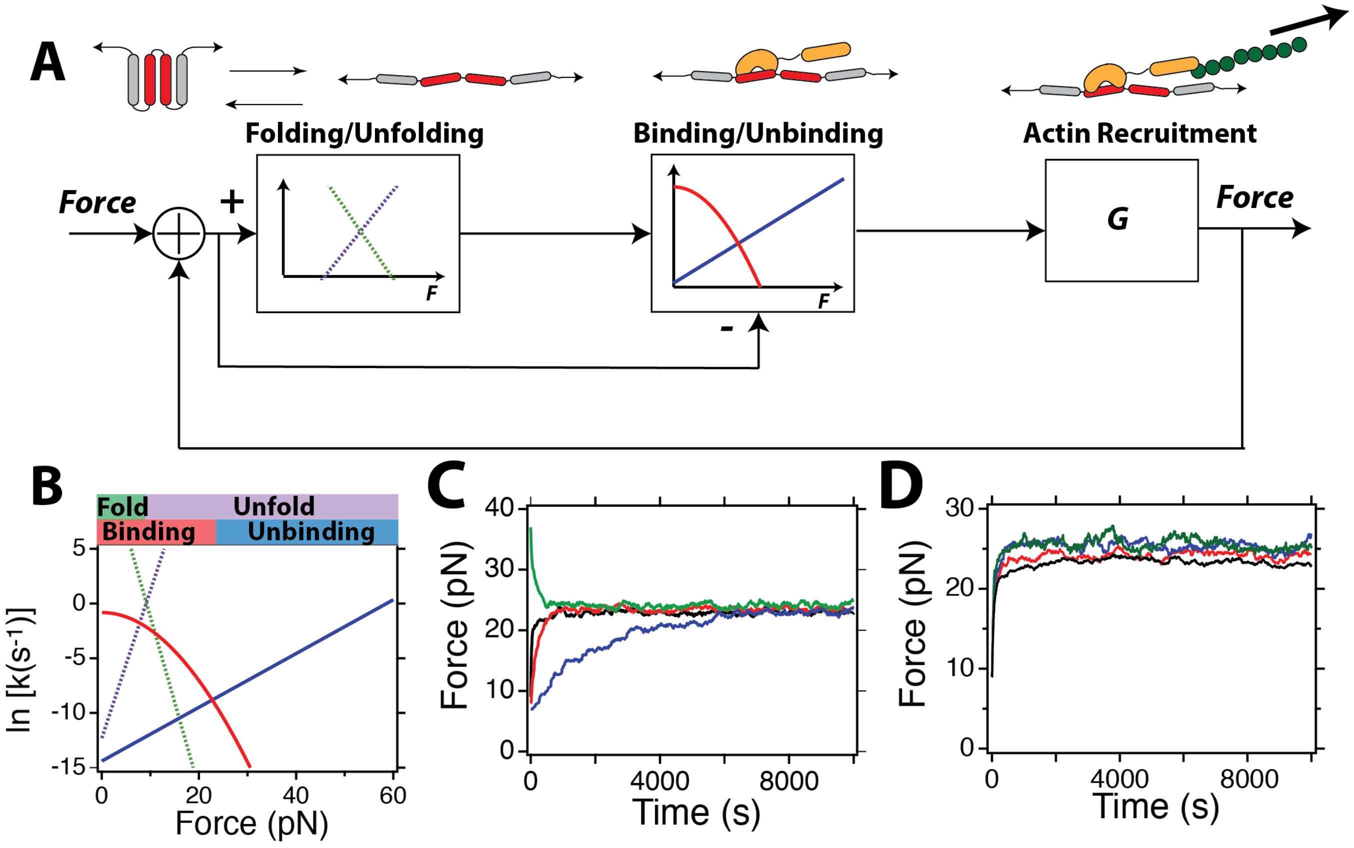

FIG. 9. **Talin-vinculin interaction defines a mechanical feedback mechanism:** (A) Block diagram for the talin-vinculin-actin linkage. (B) Folding/unfolding talin rates (green, purple), and binding/unbinding vinculin rates (red, blue), as measured experimentally. (C) Monte Carlo simulations on the talin-vinculin mechanical control system. The system operates as a negative feedback mechanism, modulating the force to maintain it constant around 23 pN. Initial forces 7, 8, 9, and 37 pN; blue, red, black, and green. (D) Robustness of the talin-vinculin-actin linkage model to the force gain introduced per bound vinculin. Force gains of 2.5 pN (black), 4 pN (red), 5 pN (blue), and 7 pN (green).

We run Monte Carlo simulations on this synthetic talin polypeptide, built by 10 of the described blocks, which have the characteristics we measured for vinculin head binding to R3 IVVI. In particular, the binding/unbinding rates of each talin module:

$$k_{\rm U} = (1.84 \times 10^{-6}) \exp{(5.54F/kT)}$$
$$k_{\rm F} = (6.42 \times 10^{6}) \exp{(-6.42F/kT)},$$

as measured in Fig. S2, where the rates are expressed in $s^{-1}$ and the distances to transition state in nm. For vinculin binding, we use:

$$k_{\rm b} = 0.0011[Vh]^2 \exp{(-\Delta W(F)/kT)}$$
$$k_{\rm u} = (5.6 \times 10^{-7}) \exp{(1.05F/kT)},$$

where $\Delta W(F) = 7.3F(\coth^{-1}(F0.11/kT) - kT/(F0.11))$, as extracted from the binding probability measurements. Figure S9B shows the force-dependence of the rates we use. At each Monte Carlo step, we evaluate the folding and binding status of each of the domains, starting with an arbitrary input force. The evolution of talin folding status is simulated from its folding and unfolding rates. If talin is unfolded, we run a Monte Carlo step with vinculin binding rates, based on the force acting on the system. If vinculin binds, the talin domain is locked on the unfolded state, and a force gain is added to the system due to actin recruitment. This force gain is 3 pN per bound vinculin, which is around the tension across vinculin-actin linkages. If vinculin is bound on a domain, we run a Monte Carlo step based on the unbinding rate.

Figure S9C shows the time evolution of the force born by talin at four different initial conditions (7, 8, 9, and 37 pN; blue, red, black, and green). Each trajectory is obtained by averaging 30 realizations. The system operates as a negative mechanical feedback system, which regulates the force to keep it around a constant value of 23 pN. This value is determined from the physics of vinculin unbinding, being the force at which vinculin binding/unbinding rates are equal. The exact setpoint force depends on the concentration of vinculin head, but just in a linear way, being the physics of the coil-to-helix transition the main determinant. At 8 nM, the equilibrium force would be 21 pN, and at 40 nM 25 pN. Additionally, the system is robust to the gain force per bound vinculin, which just controls the kinetics of equilibration. Figure S9D shows simulations using gains of 2.5, 4, 5, and 7 pN, and all equilibrate to the same equilibrium force.

### XIII. METHODS

#### A. Magnetic tweezers setup

All the experiments were done on our custom-made magnetic tweezers setup, as described before [26, 27]. The instrument is built on top of an inverted microscope (Olympus IX- 71/Zeiss Axiovert S100) using a 63X or a 100X oil-immersion objective (Zeiss/Olympus), mounted on a nanofocusing piezo actuator (P-725; Physik Instrumente). The fluid chambers are illuminated using a collimated cold white LED (ThorLabs). Images were acquired using a CMOS Ximea MQ013MG-ON camera. Molecules were tethered to superparamagnetic Dynabeads M-270 beads (2.8 $\mu$m diameter). Calibrated force were applied using either a voice-coil mounted pair of permanent magnets (Equipment Solutions), or a magnetic tape head (BRUSH 902836). Both the voice-coil and magnetic head were maintained under electronic feedback using a home-made PID circuit controller. The force was calibrated as described previously [26, 27]. The data acquisition and control of the piezo was done using a multifunction DAQ card (NI USB-6289, National Instruments).

#### B. Image processing

Image processing was done by custom-written software written in C++/Qt, fully available on the lab's website (http://zeptowatt.com), and as described before [26].

#### C. Fluid chamber preparation

All experiments are done in custom-made fluid chambers built by two sandwiched glass coverslips (Ted Pella), separated by a laser-cut parafilm pattern. The bottom glass surfaces are silanized as described before [26]. The top glass is treated with Repel Silane (Sigma-Aldrich) for hydrophobization. The fluid chamber is sealed by sandwiching both surfaces with the parafilm pattern at 80 C. The fluid chambers are functionalized to add the HaloTag ligand and reference beads as described before [26]. Both the fluid chambers and magnetic beads are passivized using TRIS blocking buffer (20 mM Tris-HCL pH 7.4, 150 mM NaCl, 2mM MgCl2, and 1% w/v/ sulfhydryl blocked-BSA).

#### D. Protein expression and purification

Polyprotein constructs were engineered using a combination of BamHI, BglII, and KpnI restriction sites in pFN18a restriction vector, as described previously [26]. Our protein construct for the magnetic tweezers experiment contains the R3 IVVI or R3 WT mouse talin domain, followed by eight I91 domains, and flanked by an N-terminal HaloTag enzyme and a C-terminal AviTag for biotinylation. For purification purposes, a (His)6-tag was also present before the AviTag. BLR (DE3) or ERL competent cells were grown at 37°C, and the protein over-expression was induced with 1mM Isopropyl -D-1-thiogalactopyranoside (IPTG, Sigma) overnight at 25°C. Cells were resuspended in 50 mM sodium phophate buffer pH 7.0, 300 mM NaCl, 10% glycerol, and lysed. The proteins were purified from the lysate first with a Ni-NTA affinity resin, followed by size exclusion chromatography using a Superdex-200 HR column in 10 mM HEPES buffer pH 7.2, 150mM NaCl, 10% v/v glycerol and 1mM EDTA. The purified proteins were then concentrated to 50-100 $\mu$M and biotinylated in 50 mM Bicine buffer pH 8.3, 10 mM magnesium acetate, 10 mM ATP, 100 $\mu$M biotin and 2.5 $\mu$g biotin ligase BirA enzyme (Avidity), at 40°C for 4 hours. Human vinculin head was expressed and purified following an analogous procedure, skipping the biotinylation process.

#### E. Single molecule measurements

All experiments were carried out in HEPES buffer (Hepes 10 mM pH 7.2, NaCl 150 mM, EDTA 1 mM) containing 10 mM ascorbic acid (pH 7.3) to avoid oxidative damage, and the desired vinculin head concentration. The protein was incubated on the fluid chamber for 30 minutes at a concentration of ∼3 nM, and then washed with measuring buffer to remove the free molecule. The beads were incubated to bind the protein for around 2-5 minutes, after which a force of 4 pN was applied. Experiments begin with a z-stack library of the magnetic and reference bead. From then, the z-position displacements are calculated in real time using our image processing method, described previously [26].

#### F. Single molecule data analysis

Our data acquisition software writes a four-column binary file containing the time mark, extension, force, and magnet position/current. We visualize and analyze the data using custom written software written in Igor Pro (Wavemetrics). All data is acquired at frequencies ranging from 1,000 up to 1,600 Hz. Data is smoothed with a 4th order Savitzky-Golay filter using a box size of N=101.

The folding/unfolding states of talin are automatically detected using a double threshold algorithm. Briefly, a length histogram is calculated on the smoothed trajectory. A double gaussian fit is done to determine the extensions of the folded ($L_F$) and unfolded ($L_U$) states. We set the threshold for the folded state at $L_F + \sigma/2$, being $\sigma$ the standard deviation of the gaussian fit, and the equivalent one for the unfolded state. From this threshold definition the extension time series are idealized by assigning each data point onto the folded or unfolded domain regarding what threshold they overcome.

Vinculin binding was detected in base of the fingerprint on talin dynamics—no folding dynamics at a force of 9 pN (R3 IVVI) or 5 pN (R3 WT). Binding times were measured as the time taken from the moment the measuring force was set—always using 4 pN (R3 IVVI) or 3 pN (R3 WT) as a reference force—to the time where the contraction that precedes the arrest of talin folding dynamics happened. At forces where talin does not refold ($>$ 10 pN), vinculin binding was detected from the binding step. In every case, a probe pulse was set at 9 pN to verify the binding status. In all our experiments, the presence of a binding step resulted in no talin folding dynamics at 9 pN (R3 IVVI) or 5 pN (R3 WT). Unbinding of vinculin was done at forces of 40, 50, and 60 pN, after which a probe pulse at 9 pN (R3 IVVI) or 5 pN (R3 WT) was set to confirm vinculin unbinding by the recovery of talin folding dynamics.
